# In Vivo Optical Clearing of Mammalian Brain

**DOI:** 10.1101/2024.09.05.611421

**Authors:** Giovanni Talei Franzesi, Ishan Gupta, Ming Hu, Kiryl Piatkevich, Murat Yildirim, Jian-Ping Zhao, Minho Eom, Seungjae Han, Demian Park, Himashi Andaraarachchi, Zhaohan Li, Jesse Greenhagen, Amirul Muhammad Islam, Parth Vashishtha, Zahid Yaqoob, Nikita Pak, Alexander D Wissner-Gross, Daniel A. Martin-Alarcon, Jonathan J. Veinot, Peter T. C. So, Uwe Kortshagen, Young-Gyu Yoon, Mriganka Sur, Edward S. Boyden

**Affiliations:** McGovern Institute, Massachusetts Institute of Technology; Cambridge, MA, U.S.A.; HHMI; Cambridge, MA, U.S.A.; Department of Biological Engineering, Massachusetts Institute of Technology; Cambridge, MA, U.S.A.; Departments of Brain and Cognitive Sciences, Massachusetts Institute of Technology; Cambridge, MA, U.S.A; Departments of Media Arts and Sciences, Massachusetts Institute of Technology; Cambridge, MA, U.S.A; Koch Institute, Massachusetts Institute of Technology; Cambridge, MA, U.S.A; K. Lisa Yang Center for Bionics, Massachusetts Institute of Technology; Cambridge, MA, U.S.A; Center for Neurobiological Engineering, Massachusetts Institute of Technology; Cambridge, MA, U.S.A; Baylor College of Medicine; Dallas, TX, U.S.A; School of Life Sciences, Westlake University; Cloud Town, China; Westlake Laboratory of Life Sciences and Biomedicine, Westlake University; Cloud Town, China; Institute of Basic Medical Sciences, Westlake University; Cloud Town, China; Westlake Institute for Advanced Study, Westlake University; Cloud Town, China; Lerner Research Institute, Cleveland Clinic; Cleveland, OH, U.S.A; Department of Electrical Engineering, Korea Advanced Institute of Science & Technology; Daedeok Innopolis, Republic of Korea; Department of Mechanical Engineering, University of Minnesota; Minneapolis, MN, U.S.A; Department of Chemistry, University of Alberta; Edmonton, Canada; Department of Mechanical Engineering, Massachusetts Institute of Technology; Cambridge, MA, U.S.A; Reified Labs; Cambridge, MA, U.S.A

## Abstract

Established methods for imaging the living mammalian brain have, to date, taken the brain’s optical properties as fixed; we here demonstrate that it is possible to modify the optical properties of the brain itself to significantly enhance at-depth imaging while preserving native physiology. Using a small amount of any of several biocompatible materials to raise the refractive index of solutions superfusing the brain prior to imaging, we could increase several-fold the signals from the deepest cells normally visible and, under both one-photon and two-photon imaging, visualize cells previously too dim to see. The enhancement was observed for both anatomical and functional fluorescent reporters across a broad range of emission wavelengths. Importantly, visual tuning properties of cortical neurons in awake mice, and electrophysiological properties of neurons assessed *ex vivo*, were not altered by this procedure.

## Introduction

Optical imaging of processes within living organisms is important throughout the biological sciences, and advances in our ability to image deep in living tissue have been key to furthering our understanding of normal functions and pathological physiology *in vivo*^1–3^.

A number of factors help shape the landscape of what is possible to image in vivo, and have consequently been the focus of efforts aimed at minimizing any detrimental effects they might have. Such factors include, for example, opaque tissues overlying the region of interest ^4–16^, refractive index mismatches between the tissue and the fluid the microscope lens is immersed in ^17–19^, and light absorption by the blood ^20^; however, at least in mammalian tissues such as the brain, the main limiting factor is thought to be light scattering within the tissue itself. Whether through inventing new kinds of microscope^21–26^, fluorophore^27–33^, or computational imaging strategy^34–37^, attempts to date to confront this limit accept the optical properties of living tissue as fixed, and aim to work around them. In contrast, until now, no method has that successfully altered the optical properties of a delicate tissue such as the brain, while preserving its normal physiology, has been reported.

Optical clearing of chemically preserved specimens^38–43^ has long been useful in studying the structure of tissues and the distribution of biomolecules of interest. To date, however, methods developed to clear preserved specimens involve harsh steps (e.g. complete removal of lipids, or saturation of tissue with high concentrations of chemicals like urea) that are not applicable to live tissue. While *in vivo* clearing of robust tissues such as bone, skin, ligament, and muscle have been described^16,44–47^, the conditions involved (e.g., very high concentrations of glycerol) have limited their use on delicate systems such as the mammalian brain.

Recently, Iijima and colleagues^48^ reported a pharmacological method aimed at decreasing scattering in the brain, putatively by ameliorating surgery-related edema and ischemia^48^ via the chronic administration of 5% glycerol to a mouse’s drinking water. Brain physiology was not characterized, however, and the enhancements reported were on the order of 25% or so.

We here show that it is possible, by acutely superfusing the cortical surface prior to imaging with aCSF supplemented with a small amount of higher RI bioinert components, to increase the signal above background from neurons at depth up to several-fold, without compromising neuron health or network activity *in vivo*.

Because most light scattering in tissue is caused by refractive index (R.I.) mismatches between higher R.I. components, such as lipid membranes^1^, and lower R.I. components, such as aqueous compartments inside and outside cells^1,49,50^, we postulated that we could reduce light scattering by increasing the R.I. of the extracellular aqueous compartment, via infusing a bioinert material with a refractive index higher than that of the extracellular fluid (∼1.33-1.34). We discovered that raising by 0.01 the refractive index of artificial cerebrospinal fluid (aCSF) used to superfuse the living mouse cortex prior to in vivo imaging, through the addition of a modest concentration of any one of several common biocompatible materials (e.g., 1.5 mM 40kDa molecular weight dextran), was sufficient to significantly improve one-photon and two-photon imaging of brain structure and dynamics.

In summary: improvements were observed throughout the volume imaged (up to the limits of one-photon and two-photon imaging with conventional hardware), with the effect of optical clearing increasing with depth. The enhancement was especially appreciable near the depth limits of an imaging technology (i.e., below 100-150µm when imaging under one photon microscopy, and below 400-500µm when imaging under two photon microscopy), where, at baseline and under control conditions, signals from cells were extremely low, or even indistinguishable from background. After clearing, many more cells became visible, and those that were identifiable at baseline showed, for the deepest range imaged (450 to 550µm) a median increase in signal above background of ∼+385%, while many cells that were barely distinguishable above background at baseline often showed order-of-magnitude improvements or better.

Live tissue optical clearing significantly increased the signal obtained from all fluorophores tested, which had peak emission wavelengths ranging from green (GFP^51^ and genetically encoded calcium sensors of the GCaMP family^29,30^, with excitation (ex) and emission (em) peaks of 488nm and 507 nm respectively), to red (tdTomato^52^, ex 554 nm/em 581 nm), to far red/NIR (iRFP682^53^ and the genetically encoded voltage indicator Archon^54,55^, ex 663 nm/em 682 nm and ex 637 nm/em 664 nm, respectively).

We assessed cell health and cell physiology under live clearing treatment using multiple standard assays, finding excellent safety and preservation of cell functionality. To evaluate whether complex properties of *in vivo* neural networks behavior and coding, which arise from the interactions of neurons within and across a brain region, were also unaffected, we compared the visual orientation tuning of neurons in the primary visual cortex of awake mice undergoing visual stimulation following superfusion with control aCSF vs. aCSF containing 1.5 mM 40 kDa dextran; no significant difference between the two conditions was found, suggesting that even network-scale physiology in vivo was preserved after optical clearing.

While we chose to focus on cheap, widely available biocompatible materials with an extensive track record of use in biological or medical applications, custom materials designed from the ground up for *in vivo* optical clearing could, in principle, offer a much greater degree of refractive index matching and, hence, efficacy, while causing minimal osmolarity increases.

Silicon is an appealing material for this application because of its high refractive index (4.5^56^, compared to 1.41-1.43 for dextran, PEG, and iodixanol^57–59)^, and low toxicity in vivo^60^. We therefore developed PEG-functionalized silicon nanoparticles, as a proof of principle nanoparticle refractive index matching material, and showed their effectiveness *ex vivo*, obtaining an improvement, on average, of over 250% in the brightness above background of target beads imaged through an acute brain slice maintained under standard electrophysiology conditions. Thus optical clearing of living tissues may be implemented in principle through many different means, opening up not only many practical applications in neuroscience in the short term, but also straightforward paths in the future for diversifying and extending this toolbox.

## Results

### Initial screening of candidate in vivo clearing reagents in brain slice

Acute mouse brain slices are convenient substrates for rapidly screening potential candidates for optical clearing of live tissue, before testing the most promising ones in vivo. A simple assay that allows a quantitative assessment of changes in tissue transparency following incubation with a reagent is imaging an array of beads of uniform brightness through a tissue slice while keeping the imaging parameters constant between baseline and post-incubation imaging sessions.

Following one hour incubation in control aCSF, properties of the tissue remained similar to baseline, with a slight decrease in transparency, possibly correlated with deteriorating health of the tissue over time, (**Figure 1A, C**; median −60.18%, of brightness over background of beads imaged through the tissue, p = 0.0201, two-sample Kolmogorov-Smirnov test, n = 51 beads before, 46 after, incubation, from 5 slices from 2 mice; see **Supp Table 1** for full statistics). In contrast, incubation with aCSF to which a reagent (dextran of 40 kDa molecular weight, PEG of 10 kDa molecular weight, or iodixanol) was added at a concentration sufficient to raise the aCSF’s refractive index by 0.01 (1.5 mM, 6mM, and 40 mM respectively), made the tissue significantly more transparent (**Figure 1B** through **H** and **S.I. Figure 1**; median +217.67, p = 6.4582*10^-29^ for baseline vs. post-incubation, two-sample Kolmogorov-Smirnov test, n = 67 beads before incubation, n = 69 beads after incubation, over 5 slices from 3 mice for dextran-aCSF; median +429.21%, p = 4.4210*10^-41^, two-sample Kolmogorov-Smirnov test, n = 82, 124 beads for before vs. after incubation, from 4 slices from 3 mice, for PEG-aCSF; median +127.63%, p = 4.3646*10^-6^, two-sample Kolmogorov-Smirnov test, n = 90, 244 for before vs. after incubation, from 7 slices from 3 mice, for iodixanol-aCSF; see **Supp. Table 1** for full statistics). Thus, multiple chemically independent molecules could, via raising the refractive index of aCSF, result in improved brain slice transparency.

**Fig. 1.**
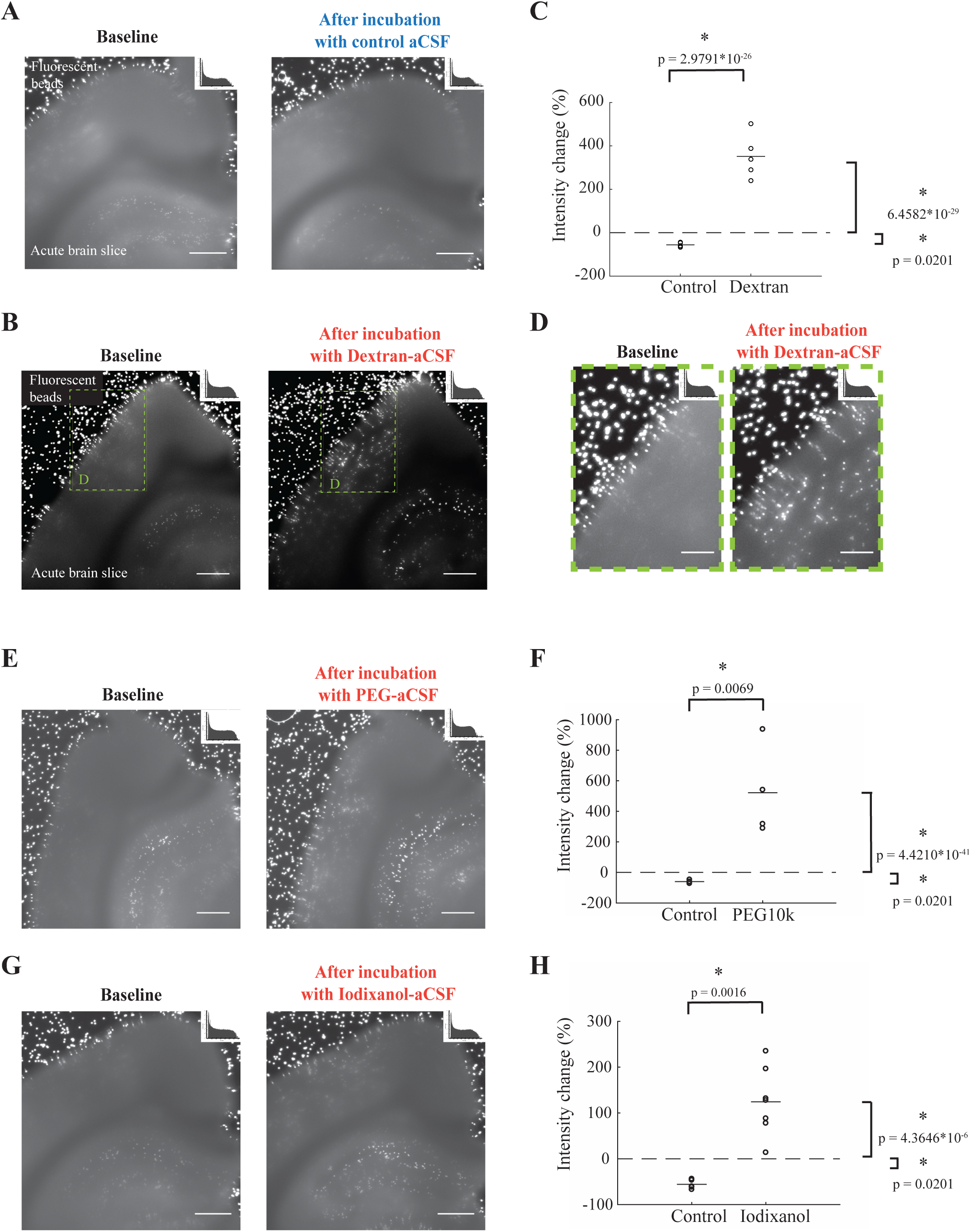
Optical clearing of acute brain slices incubated for one hour in oxygenated higher-refractive index aCSF, assessed by transmission imaging of fluorescent beads. A. Array of fluorescent (emission 645nm) 15μm-diameter polystyrene beads imaged though a 250μm-thick acute brain slice imaged with identical illumination and acquisition parameters before (left) and after (right) incubation for 1hr in plain aCSF under standard conditions for *ex vivo* electrophysiology (see Methods for details). Display settings are the same for both conditions. Scale bar = 500µm. The histogram for the raw pixel values is shown, plotted in semilogarithmic form, in the top right corner of each image. B. as in A., for incubation for 1hr in aCSF containing 1.5mM Dextran 40kDa and having a refractive index higher by 0.01 compared with plain aCSF (Dextran-aCSF). Scale bar = 500µm. C. Change in measured intensity of the signal from the beads imaged through slices incubated in either standard aCSF (control) or Dextran-aCSF. Each circle represents the average value for beads imaged through a slice. Solid line segments indicate means, the dashed line marks 0 (no change). D. Sub-region from the slice in B., shown at higher magnification. Scale bar = 150µm. The histogram for the raw pixel values is shown, plotted in semilogarithmic form, in the top right corner of each image. E. and G. Same as B., for aCSF containing 6mM PEG 10kDa and 40mM Iodixanol, respectively. F. and H. same as C., for aCSF containing 6mM PEG 10kDa and 40mM Iodixanol, respectively. In C., F., and H., an asterisk denotes statistical significance at the 0.05 level. See Results and Supp. **Table 1** for full statistics. [e.g., (A)]. Additional callouts are indicated the same way, but without the bold format.

Thus, multiple chemically unrelated molecules could, via raising the refractive index of aCSF, result in increased brain slice transparency.

### Initial in vitro and ex vivo safety assessment of candidate live tissue clearing reagents

As an initial safety assessment of the candidate reagents for in vivo optical clearing, we characterized the electrophysiological properties of mouse hippocampal neurons in culture by standard patch clamp methods, for neurons incubated for 1hr in standard Tyrode solution vs. Tyrode containing one of the three reagents tested, at the same concentration as used for live tissue optical clearing. For all three reagents tested, none of the electrophysiological properties measured, i.e. resting membrane potential, input resistance, cell membrane capacitance, membrane time constant, spike threshold, and spike duration, showed any significant change (**S.I. Figure 2** p > 0.1 for all conditions and parameters, two-sample Kolmogorov-Smirnov test; n = 12 neurons from 5 cultures, control, n = 11 neurons from the same 5 cultures, treated, for the dextran case; n = 17 control cells, n = 15 treated cells, from one culture, for the PEG case; n = 15 neurons from 3 cultures, control, n = 14 neurons from 2 cultures, treated, for the iodixanol case; see **S.I. Figure 2** for full statistics).

**Fig. 2.**
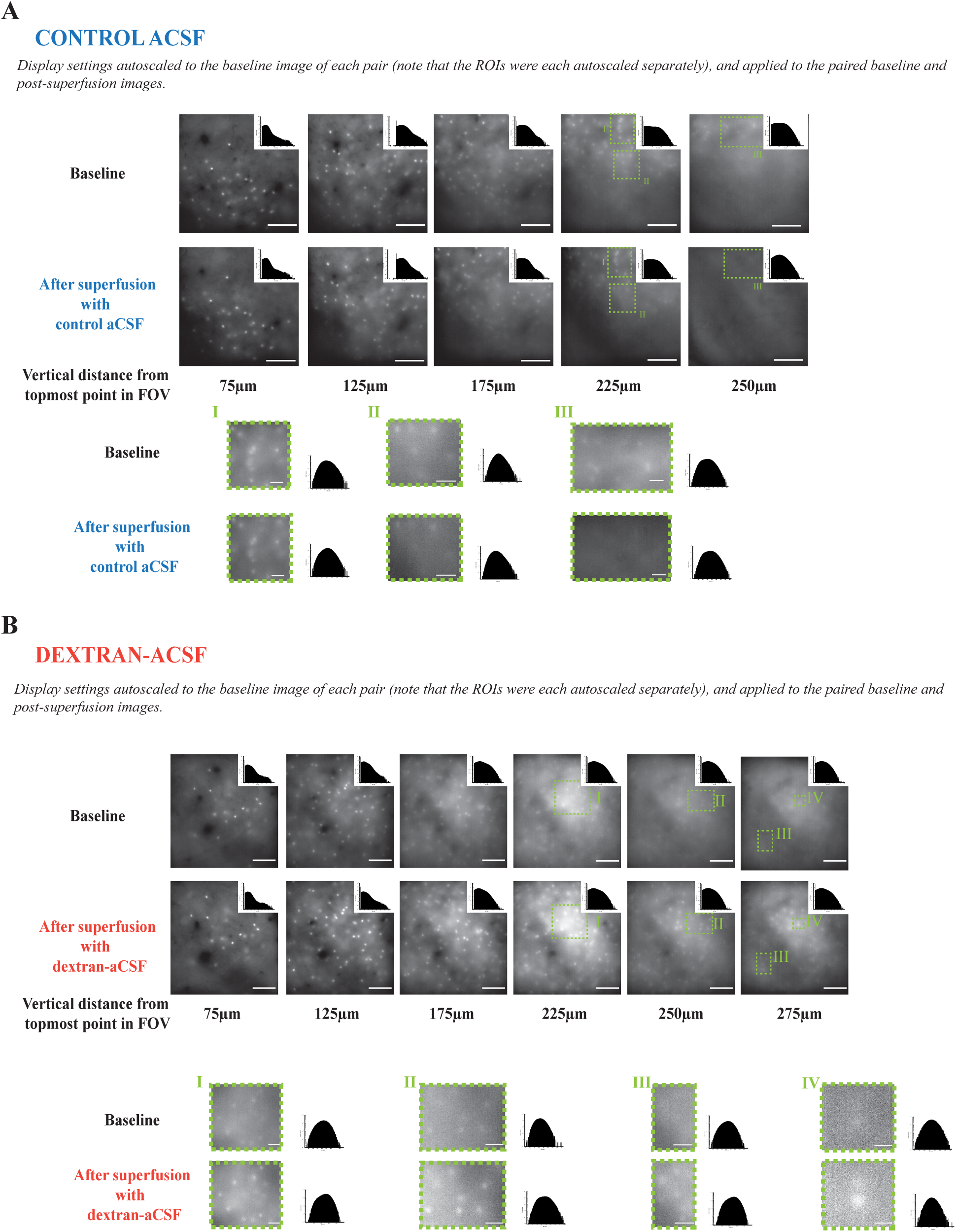
Superfusion of the cortical surface for one hour with the modified aCSF enhances imaging at depth under one photon microscopy. A. Representative maximum intensity projections, after background subtraction (see Results and Methods for details), for different depths, of tdTomato-labeled PV+ neurons in mouse somatosensory cortex, before (top) and after (bottom) 1hr cortical superfusion with control aCSF. Each maximum intensity projection is taken over 7 slices, acquired at 1.5 µm intervals, to compensate for slight misalignments in depth between the two conditions. For both before and after superfusion conditions, the image stacks were acquired under plain aCSF, using identical parameters. The settings for image acquisition and display are the same for the two conditions. Scale bar = 100µm. The highlighted regions of interest (ROI’s) I, II, and III are shown at higher magnification at the bottom of the panel. Scale bar = 25µm. Histograms for the raw pixel values are shown, plotted in semilogarithmic form, in the top right corner of the overall field of view, and to the right of the selected ROIs. The images are displayed with the brightness, contrast, minimum and maximum values determined by autoscaling in Fiji (see Methods) for the baseline image of each pair. See S.I Figure 5A. for the same images shown with settings determined by autoscaling in Fiji for the post-superfusion image. The inset in each panel shows the histogram for the raw image, plotted on a semilogarithmic scale. B. As in A., for an animal superfused for 1hr with dextran-aCSF instead of control aCSF. For the full-frame images, scale bar = 100µm. For the highlighted regions of interest (ROI’s), shown at a larger scale on the right: in I, II, and IV scale bar = 25µm; in III scale bar = 10µm. See S.I Figure 5B. for the same images shown with settings determined by autoscaling in Fiji for the post-superfusion image.

Of these three reagents, dextran 40kDa required the smallest increase in osmolarity for a 0.01 refractive index increase, so we chose to focus on it for most of our subsequent experiments. Since more than one hour might be required for diffusion of a clearing agent into deeper layers of the cortex, we repeated the cultured neuron assessments for dextran-aCSF, using a 2 hour incubation period, again without observing any significant change in electrophysiological properties (**S.I. Figure 3**; p > 0.1 for all properties assessed, two-sample Kolmogorov-Smirnov test; n = 19 neurons from 3 cultures, control, n = 12 neurons from the same 3 cultures as for control, treated; see figure for full statistics), as well as on acute mouse brain slices, again incubating for 2 hours (**S.I. Figure 4**; p > 0.1 for all properties assessed, n = 12 cells from 5 slices from 5 mice, control, n = 11 cells from the same 5 slices, treated; see figure for full statistics).

**Fig. 3.**
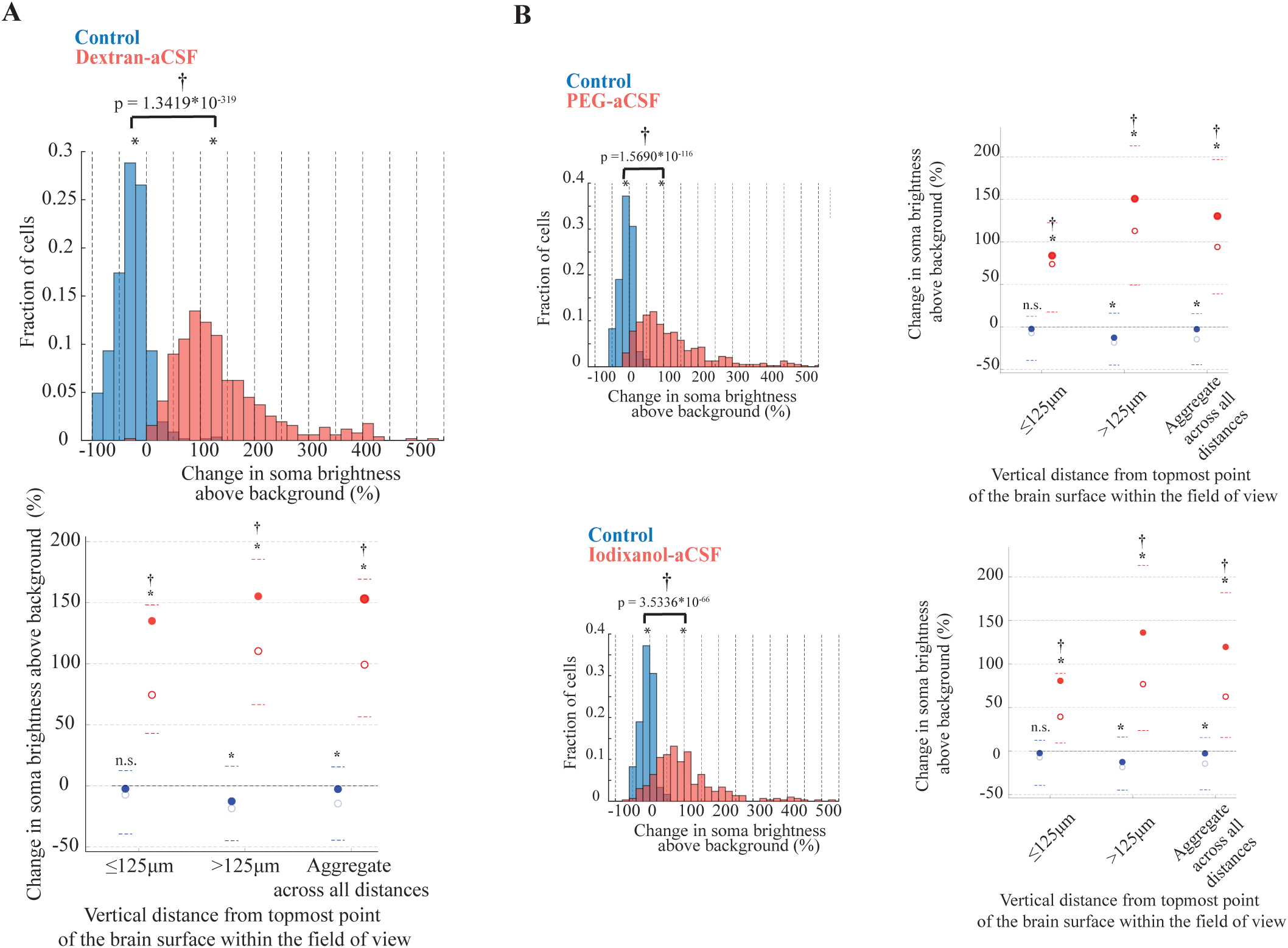
Superfusion of the cortical surface for one hou r with the modified aCSF is sufficient to, on average, more than double the intensity of the signal over background from fluorescent cortical neurons in vivo under one-photon imaging. **A.** Changes in soma brightness above background (see **Methods** for details) after superfusion vs. at baseline for cells from animals superfused with either with control aCSF (in blue) or with dextran-aCSF (in red). **Top**: histogram of such values for all cells. **Bottom:** Average (filled circles), median (empty circles), 20^th^ and 80^th^ percentiles (dashed lines in color) values for cells at depths less than 125µm from the topmost point of the brain surface, for cells deeper than 125µm from the topmost point of the brain surface, and for all cells. An asterisk denotes statistical significance at the 5% level (p <0.05) compared to baseline, † indicates statistical significance at the 5% level compared to control, n.s. = not significant at the 5% level. Across all depths, n = 1321 neurons from 4 mice for dextran-aCSF, n = 807 neurons in 3 mice for control. Reported p-values for the difference between the changes in cell soma brightness above background seen after incubation with each clearing agent vs. control were calculated with the two-sample Kolmogorov-Smirnov test. See **Results** and **Supp. Table 2** for full statistics **B.** Changes in soma brightness above background before and after superfusion either with control solutions (in blue) or with one of two modified aCSF formulations that, like Dextran-aCSF also have a refractive index (R.I.) higher by 0.01 than plain aCSF. **Top**: PEG-aCSF vs. osmolarity-matched sucrose-aCSF **Bottom**: Iodixanol-aCSF vs. osmolarity-matched sucrose-aCSF. For each, **left:** histogram of the changes in soma brightness above background (see **Methods** for details) before and after superfusion either with control solutions (in blue) or with the higher R.I. aCSF (red). **Right:** changes in soma brightness above background after superfusion for cells at depths less than 125µm from the topmost point of the brain surface, for cells deeper than 125µm from the topmost point of the brain surface, and for all cells. Filled circles indicate the average value, empty circles the median, the dotted lines indicate the 20^th^ and 80^th^ percentiles. An asterisk denotes statistical significance at the 0.1% level (p <0.001) compared to baseline, † indicates statistical significance at the 0.1% level compared to control, n.s. = not significant at the 5% level. Reported p-values for the difference between the changes in cell soma brightness above background seen after incubation with each clearing agent vs. control were calculated with the two-sample Kolmogorov-Smirnov test. For PEG10kDa- and Iodixanol-aCSF, 401 cells from 6 mice, and 297 cells from 9 mice, respectively, while for the respective controls, n = 122 cells from 2 mice, and 204 cells from 2 mice. See **Results** and **Supp. Table 3** for full statistics and details.

**Fig. 4.**
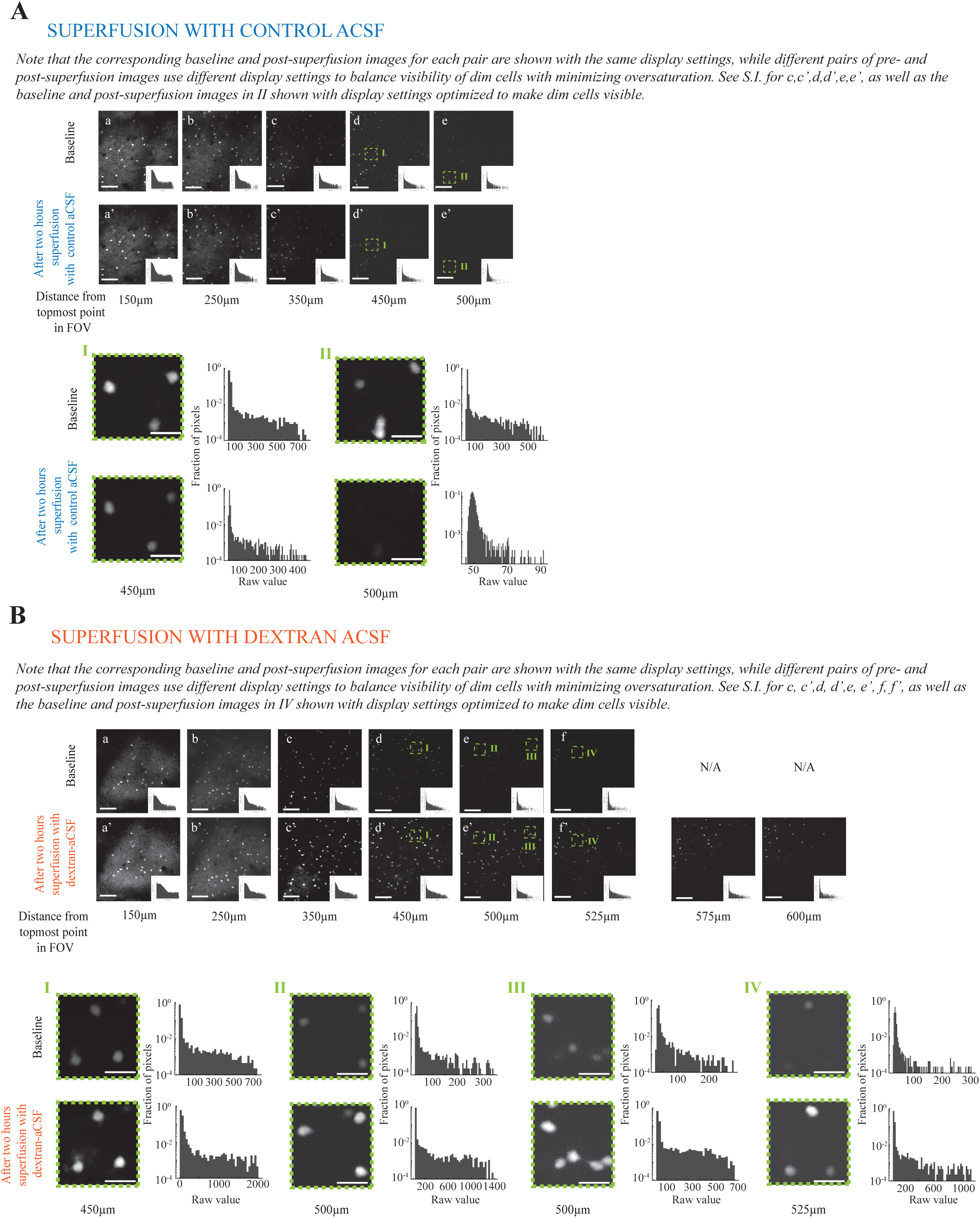
Optical clearing at depth observed with two photon imaging in vivo of fluorescently labeled primary visual cortex neurons. **A.** Representative maximum intensity projections, each from 3 imaging slices taken at 2.5µm intervals, from different depths of tdTomato labeled PV+ neurons in mouse primary visual cortex before (top) and after (bottom) 2hrs cortical superfusion with control aCSF. Scale bar = 150µm Top: before superfusion with control aCSF. Bottom: after 2hrs superfusion. Acquisition and display settings are the same for the two conditions. The images, denoised as described in **Methods,** are displayed with identical brightness, contrast, minimum and maximum values for each corresponding pair of baseline and post-superfusion images. To balance visibility of dim cells with limited oversaturation of brighter ones, such settings are adjusted between pairs of images taken at different depths. In the top set of images the full field of view is shown, while below them the enlargements of two highlighted ROI’s (**I** and **II**) are shown, together with the corresponding histograms of raw pixel values, plotted on a semilogarithmic scale. Scale bar = 37.5µm. See **S.I** Figure 6 for the same images shown with settings chosen to maximize visibility of dim cells. **B.** As in A., for a representative animal superfused with dextran-aCSF. For the image set showing the full field of view, scale bar = 150µm. For ROIs I-IV scale bar = 37.5µm. For a quantitative comparison of brightness changes following incubation please see Figure 5, **Results**, and **Supp. Table 3**.

While ex vivo and in vitro tests allow detailed characterization of electrophysiological properties of individual neurons, they do not recapitulate the complexity and activity patterns of intact brain, and may not reveal subtle effects that only appear at the network level. Thus (see below) we subsequently tested the effect of superfusion of dextran-aCSF on the orientation-specific responses of primary visual cortex neurons in awake mice. But before we discuss how dynamic imaging improves with in vivo clearing, we first discuss how we used static fluorescence in vivo to quantitatively gauge the amount of brain clearing.

### In vivo cortical clearing under one photon imaging

We imaged mouse cortical interneurons (parvalbumin (PV)-positive) expressing a static fluorophore, such as the red fluorophore tdTomato or the far red fluorophore iRFP682. One-photon microscopy is popular for in vivo imaging because of its low cost, flexibility, and potential for high imaging rates, but is highly susceptible to the effects of light scattering within tissue, and its ability to resolve features at a depth greater than 100-150µm^61^ is limited. As a measure robust to cell-to-cell fluorophore expression variability, and challenges in measuring absolute depth exactly due to brain curvature and movement, we analyzed changes in signal above background from cell bodies before vs. after superfusion, a conservative metric (because it would omit cells that were invisible or too dim at baseline to be matched with a cell that became visible post-clearing).

Imaging tdTomato-expressing neurons in the mouse visual or somatosensory cortex in vivo, after one hour cortical superfusion with plain aCSF, we observed a small decrease in imaging quality (see **Figure 2A** and **S.I. Figure 5A** for representative images, and see below for quantification of the changes in somatic brightness above background before and after superfusion). In contrast, superfusion with dextran-aCSF led to significant increases in cell body brightness above background, across the entire depth of the imaging stack, which was generally approximately 250-300µm thick overall, because of the curvature of the brain over the field of view (see **Figure 2B** and **S.I. Figure 5B** for representative images, and see below for quantification of the changes in somatic brightness above background before and after superfusion).

**Fig. 5.**
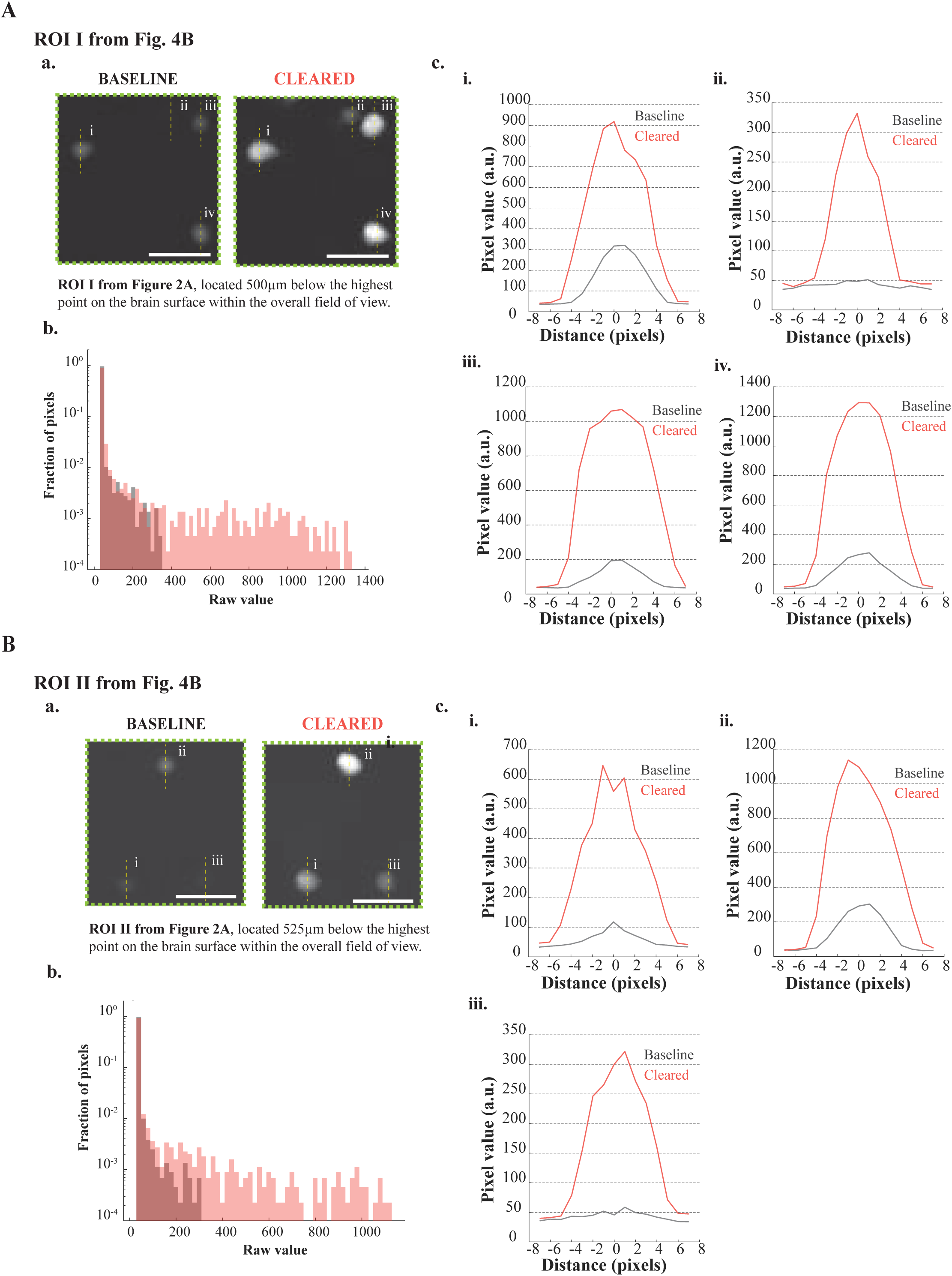
Comparison of cells in the representative ROIs highlighted in Fig. 4B, imaged before and after in vivo optical clearing. **A.** ROI I from Fig. 4B. In **a**, images shown at baseline (**left**) and after clearing (**right**). **b**: histogram of raw pixel values at baseline (**gray**), and after clearing (**red)**. **c**: profiles of raw pixel values for vertical lines passing through the estimated centroid of the cells, as indicated in a. (**solid**), or offset by 1 pixel in either direction (**dashed**), to account for potential misalignments. Note that the cells that were dimmest and least clearly identifiable at baseline, and which show the largest improvements, were excluded from the analysis described in Figure 6 below, to avoid the risk of including as part of the cell soma pixels that laid, in fact, outside of it. The estimates given in Figure 6 are therefore highly conservative ones. When plotting line profiles, in contrast, that concern is much less. **B**. As in A, for ROI II from Figure 4B.

Within the first 125µm from the topmost point on the brain surface within the area imaged (hereafter called “approximate depth”), after one hour superfusion with control aCSF, as per standard practice, we observed a small decrease in the brightness above background of cell bodies (**Figure 3A**; median −8.19%, n = 198 cells from 3 mice, p = 2.0276*10^-7^, Wilcoxon signed rank test; see **Supp. Table 2** for full statistics). Superfusion with dextran-aCSF yielded, over the same depth range, a significant increase in somatic brightness above background (**Figure 3A**; median +74.48%, n = 411 cells from 4 mice, p = 3.6842*10^-68^ for post-superfusion vs. baseline comparison, Wilcoxon signed-rank test; p = 2.2619*10^-83^ for control aCSF vs dextran-aCSF comparison, two-sample Kolmogorov-Smirnov test; see **Supp. Table 2** for full statistics). For cells below 125µm, superfusion with control aCSF caused a slight but significant decrease in soma brightness above background (**Figure 3A**, median –18.48%, n = 605 cells from 3 mice, p = 1.4976*10^-27^;Wilcoxon signed rank test; see **Supp. Table 2** for full statistics). In contrast, brightness above background of cell bodies at a distance from the topmost point on the brain surface >125µm increased after dextran-aCSF treatment (**Figure 3A**, median +110.59%, n = 874 cells from 4 mice, p = 3.22793*10^-144^ for post-superfusion vs. baseline, Wilcoxon signed-rank test; p = 6.7683*10^-245^ for control aCSF vs dextran-aCSF comparison, two-sample Kolmogorov-Smirnov test; see **Supp. Table 2** for full statistics). Aggregating over all distances from the topmost point on the brain surface within the field of view, similar results were observed (**Figure 3A**; for control aCSF median −14.82%, n = 803 cells from 3 mice, p = 1.359810^-31^, Wilcoxon signed rank test; for dextran-aCSF: median +99.22%, n = 1285 cells from 4 mice, p = 2.5189*10^-208^ for post-superfusion vs. control, Wilcoxon signed-rank test; p = 1.3419*10^-319^, dextran-aCSF vs control aCSF, two-sample Kolmogorov-Smirnov test; see **Supp. Table 2** for full statistics).

Similar results held for PEG-aCSF and iodixanol-aCSF, imaging pyramidal cells or PV+ neurons expressing the far-red fluorescent protein iRFP682^53^ in primary somatosensory cortex (**Figure 3B**; see **Supp. Table 3** for full statistics).

### In vivo cortical clearing under two-photon imaging

We next asked if in vivo clearing could boost the performance of two-photon microscopy, which allows deeper imaging than one photon microscopy, and thus is commonly used for in vivo functional brain imaging. We imaged PV+ neurons, expressing tdTomato, in mouse primary visual cortex (V1), before and after a 2-hour superfusion (the longer duration allowed for diffusion to the deeper layers of the cortex accessible by two-photon microscopy) with control vs. high refractive index aCSF. While after superfusion with control aCSF the cells were generally less visible (**Figure 4A**, **S.I. Figure 6A**, and **S.I. Figure 7A**; see below for quantification of the changes in somatic brightness above background before and after superfusion), superfusion with dextran-aCSF significantly improved imaging quality (**Figure 4B**, **Figure 5, S.I.** Figure 6B and **S.I. Figure 7B**; see below for quantification of the changes in somatic brightness above background before and after superfusion).

**Fig. 6.**
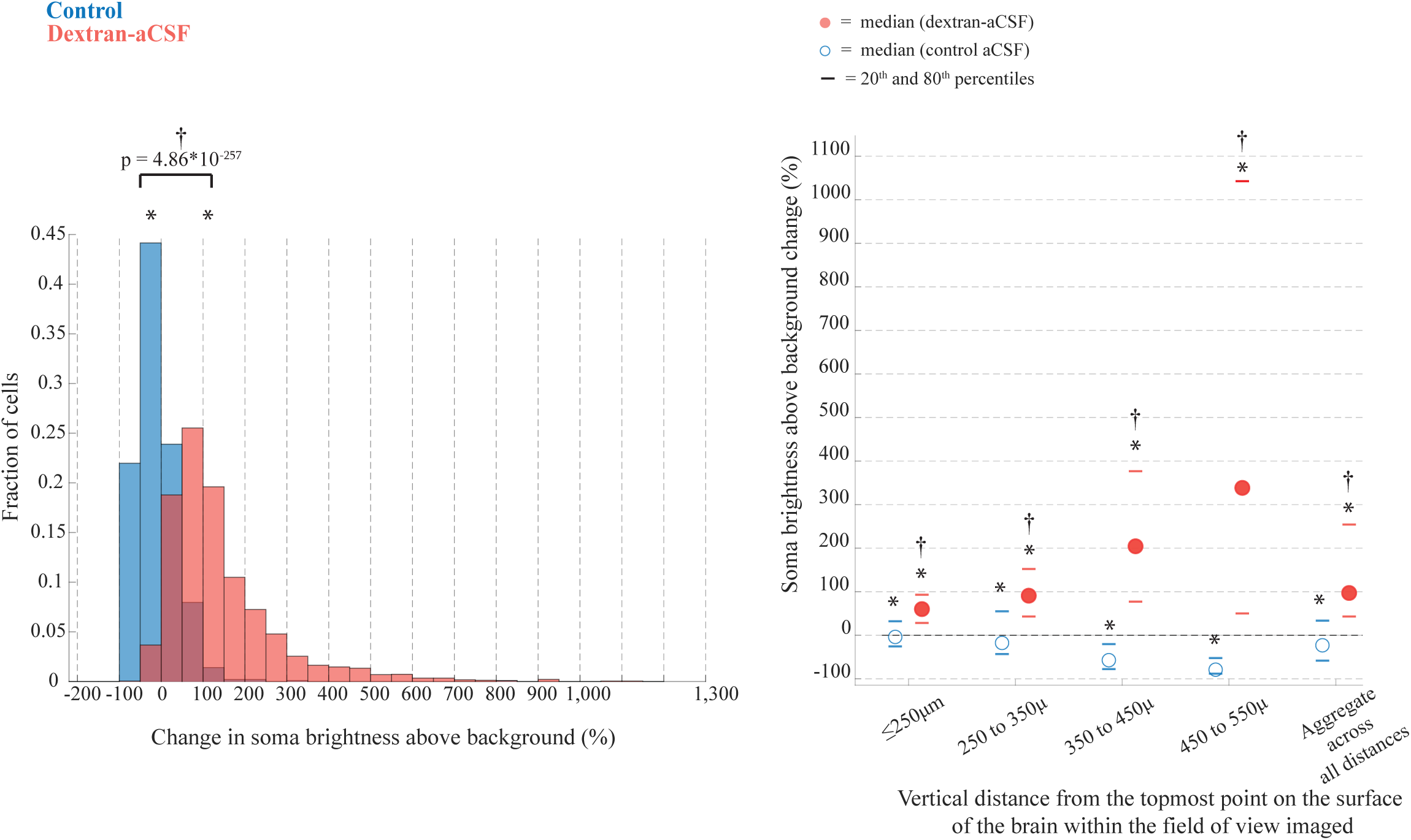
Superfusion with dextran-aCSF, on average, more than doubles the intensity of the signal over background from fluorescent cortical neurons in vivo imaged under two-photon microscopy. Changes in soma brightness above background (see **Methods** for details) after superfusion vs. at baseline for cells from animals superfused with either with control aCSF (in blue) or with dextran-aCSF (in red). **Top**: histogram of such values for all cells. **Right:** Average (filled circles), median (empty circles), 20^th^ and 80^th^ percentiles (dashed lines in color) values for cells at depths less than 250µm from the topmost point of the brain surface, for cells located between 250 and 350µm, between 350 and 450µm, between 450 and 550µm from the topmost point of the brain surface, and for all cells, regardless of depth. An asterisk denotes statistical significance at the 5% level (p <0.05) compared to baseline, † indicates statistical significance at the 5% level compared to control. P-values for the difference between the changes in cell soma brightness above background seen after incubation with each clearing agent vs. control were calculated with the two-sample Kolmogorov-Smirnov test. See **Results, Supp. Table 4**, and **Supp. Table 5** for full statistics.

**Fig. 7.**
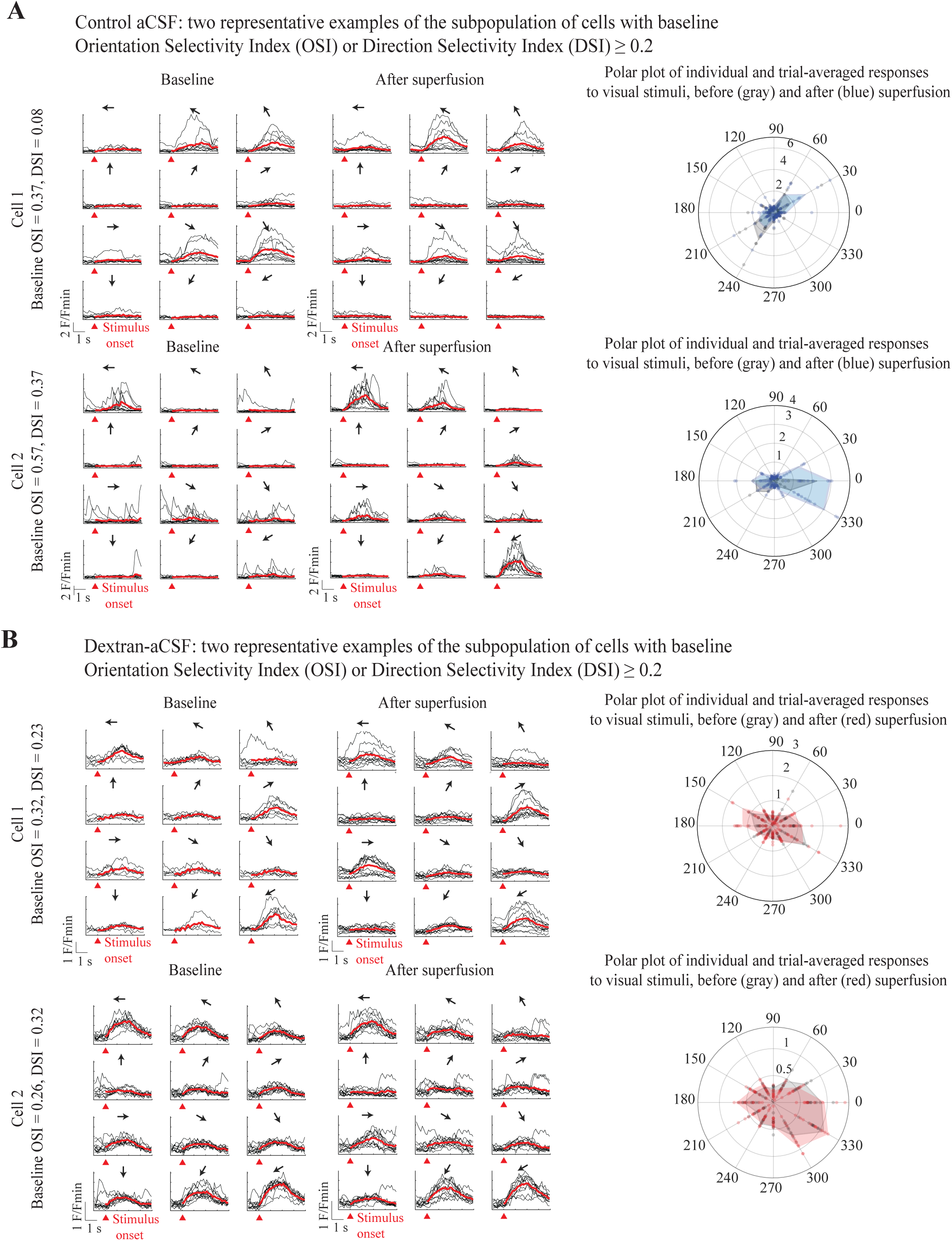
Visual tuning is preserved following two hours superfusion with dextran-aCSF – representative neurons from animals superfused with control aCSF or dextran-aCSF. **A.** Two representative neurons among those with baseline Orientation Selectivity Index (OSI) or Direction Selectivity Index (DSI) ≥ 0.2. Left and center: raw (black) and average (red) traces aligned to the time of visual stimulation for each of the directions of drifting gratings employed. The black arrow indicates the direction of the stimulus. Right: polar plots of the response to stimuli with different directions for individual trials (filled circles) and averaged across all trials (shaded area). Before superfusion is shown in gray, after superfusion in blue. See Methods for details**. B.** as in A., for representative cells with OSI or DSI ≥ 0.2 from animals superfused with dextran-aCSF. In the polar plots, baseline responses are shown in gray, responses after superfusion in red. Changes in the preferred orientation (PO) of visually tuned cells following superfusion, for control aCSF (blue) vs. dextran-aCSF (red) treated animals

For neurons located above approximately 250µm from the topmost point of the brain surface within the field of view, after two hours superfusion with control aCSF we observed a slight reduction in cell body brightness above background (**Figure 6**; median −3.8063%, n = 322 cells from 2 mice, p = 1.4182*10^-6^, Wilcoxon signed rank test. See **Supp. Table 4**, **5** for full statistics). For neurons in this depth range, superfusion with dextran-aCSF yielded, in contrast, a significant increase in brightness (**Figure 6**; median +60.09%, n = 648 cells from 3 mice, p ∼ 0 (value too small to be computed by MATLAB’s native function, henceforth “0*”) for post-superfusion vs. baseline, Wilcoxon signed-rank test; See **Supp. Table 4**, **5** for full statistics). As expected, the effect of optical clearing became progressively more pronounced for cells located at increasing depth, the median increase being +90.86%, +204.3%, and +338.42% for cells 250 to 350, 350 to 450, or 450 to 550µm from the topmost point of the brain surface within the field of view, respectively (n = 697, 728, 243 cells from 3 mice, p = 0*, 0*, 0*, 1.4868*10^-38^ for baseline vs. post-superfusion comparison, respectively, Wilcoxon signed-rank test). Within the same depth ranges, the corresponding values for animals that had been superfused with control aCSF were −17.60%, −57.20%, and −79.93% (n = 343, 385, and 42 cells from 2 mice, p = 1.911810^-14^, 3.2608*10^-40^, 1.1095*10^-07^ for baseline vs. post-superfusion comparison, respectively, Wilcoxon signed-rank test). All comparisons between cells from animals superfused with dextran-aCSF vs. control aCSF were highly significant (p = 2.5855*10^-66^, 5.6465*10^-67^, 7.1331*10^-118^, 5.1367*10^-26^; two-sample Kolmogorov-Smirnov test.

Considering all cells together, regardless of the depths from the topmost point of the brain surface within the field of view they were located at, superfusion with control aCSF led to a small reduction in somatic brightness above background (median = −23.04%, n = 992 cells from 2 mice, p = 0*, Wilcoxon signed-rank test), in contrast to superfusion with high refractive index aCSF, which significantly increased somatic brightness above background (median = +97.08%, n = 2316 cells from 3 mice, p-value for baseline vs. post-superfusion = 0*, Wilcoxon signed-rank test; p-value for dextran-aCSF vs. control aCSF superfusion = 4.8566*10^-257^, two-sample Kolmogorov-Smirnov test). See **Supp. Tables 4** and **5** for full statistics.

### Functional imaging of visual responses in awake mice following optical clearing

Since dextran-ACSF did not disrupt common electrophysiological measures of neural function in vitro, we next probed whether dextran-ACSF altered neural coding properties observed at the network level, in vivo. We imaged activity from primary visual cortex neurons (layer II/III) expressing a genetically encoded calcium sensor (see **Methods** for details) in awake headfixed mice undergoing visual stimulation and compared the visual response properties of the cells before vs. after 2 hours of superfusion with either control aCSF or dextran-aCSF. As a metric of neural coding robust to normal experimental variability^62–64^, we focused on the preferred orientation of visually tuned cells^65–67^. In both animals superfused for 2hrs with control aCSF and animals superfused with dextran-aCSF, cells showed similar activity patterns in response to visual stimulation before and after superfusion (see **Figure 7A, B** and **S.I. Figure 8** for representative examples).

**Fig. 8.**
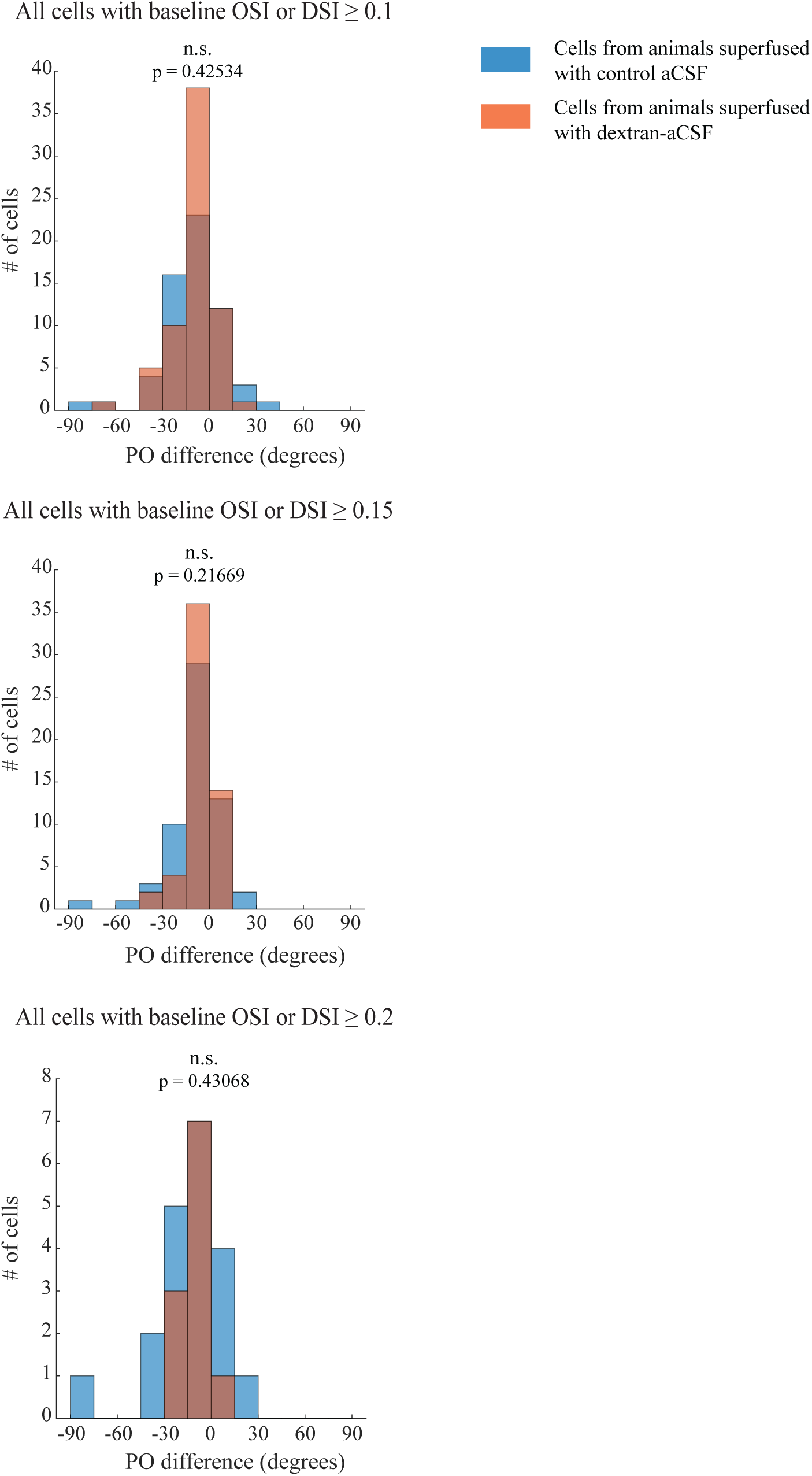
Visual tuning is preserved following two hours superfusion with dextran-aCSF – population data for neurons having baseline orientation or direction selectivity indices above 0.1, 0.15, or 0.2. Histograms showing the distribution of changes in preferred orientation (PO) before vs. after superfusion for all cells above a given threshold OSI or DSI value. From top to bottom, the threshold employed was 0.1, 0.15, 0.2. See **Results** for full statistics. Values for cells from animals superfused with control aCSF are displayed in blue, those for animals superfused with dextran-aCSF in red. The p-value above each pair of histograms is for the comparison between PO changes in cells from animals superfused with control aCSF versus those observed in cells from animals superfused with dextran-aCSF (Kolmogorov-Smirnov two-tailed test, see **Results** for full statistics).

Orientation selectivity index (OSI) and direction selectivity index (DSI) are normalized metrics for characterizing the responses of primary visual cortex cells to stimuli that are commonly used^22,68,69^, often in conjunction with the classic drifting grating stimuli. The former metric quantifies how much more likely a cell is to fire to, in our case, gratings having a certain angle, while the latter takes into account both the gratings angle and the direction in which the gratings are moving (e.g. gratings at 30° that move from the top of the screen to the bottom are different from gratings that move from the bottom towards the top).

Comparing the changes in preferred orientation that took place following superfusion in cells from control animals and animals treated with dextran-aCSF, there was no statistically significant difference for visually tuned cells that had a baseline OSI or DSI^22,68,69^ thresholded at 0.1, 0.15, or 0.2 (**Figure 8**, p = 0.42534, 0.21669, 0.43068, respectively, two-sample Kolmogorov-Smirnov test, n = 47, 20, 12 cells, out of a total of 62 cells that could be confidently matched before and after superfusion, from 2 control animals, and 52, 31, 13 cells out of a total of 67 cells that could be confidently matched before and after superfusion, from 3 dextran-aCSF treated animals, respectively). Histograms showing the distribution of OSI and DSI values before and after superfusion, for control and dextran-aCSF treated animals, are shown in **S.I. Figure 9**. A quantitative comparison of the differences in OSI and DSI changes between baseline and after superfusion comparing control and dextran-aCSF treated animals is shown in **S.I. Figure 10**, and reveals no statistically significant differences between control and dextran-aCSF treated animals, regardless of whether all cells were included, or only cells with OSI or DSI baseline values above a certain threshold, e.g. 0.1 and 0.15 (for OSI changes, p = 0.2474, p=0.4894, p = 0.1238; for DSI changes, p = 0.1185, p=0. 6753, p = 0.5378, two-sample Kolmogorov-Smirnov test, n = 62, 47, 20 cells from 2 control animals, n = 67, 52, 31 cells from 3 dextran-aCSF treated animals, respectively). Looking at OSI and DSI combined, i.e., categorizing cells as visually tuned if either index met a certain criterion, confirmed that no cells which showed no evidence of visual tuning at baseline (selectivity index < 0.05) became strongly tuned (selectivity index ≥ 0.15) after superfusion, nor did any cell exhibiting strong tuning fall in the lowest-tuning category after superfusion (see **S.I. Figure 11**, **Supp. Table 6**, and **Supp. Table 7**).

**Fig. 9.**
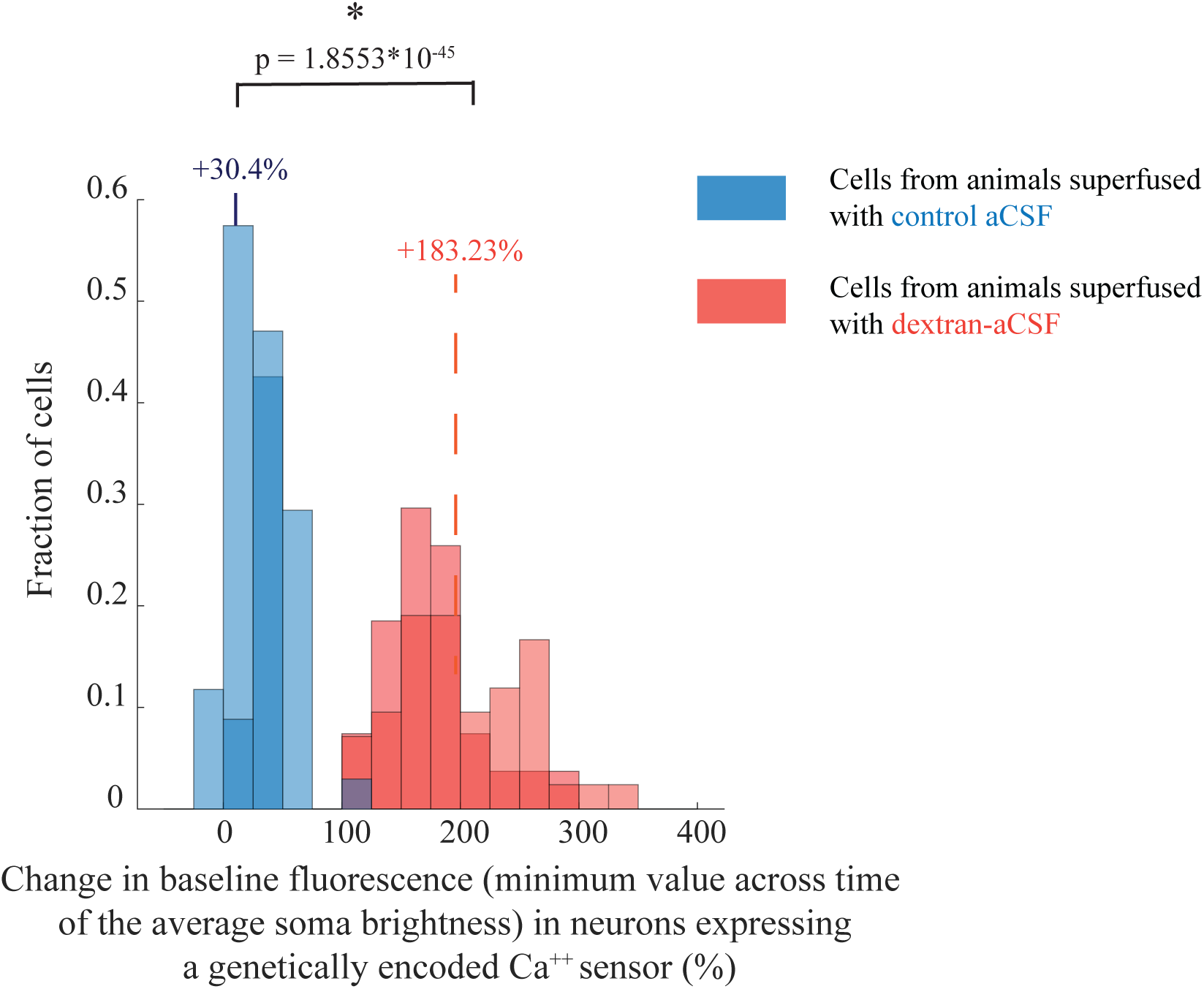
Optical clearing at depth observed under two-photon imaging of labeled visual cortex neurons expressing a genetically encoded calcium indicator. Histograms for changes after two hours cortical superfusion in baseline fluorescence (calculated as the minimum over time of average soma brightness above background) of neurons expressing a genetically encoded Ca^++^ indicator. Values for cells from animals superfused with control aCSF are displayed in blue, while those for cells from animals superfused with dextran-aCSF are shown in red (see **Methods** for details, **Results** and **Supp. Table 8** for full statistics).

**Fig. 10.**
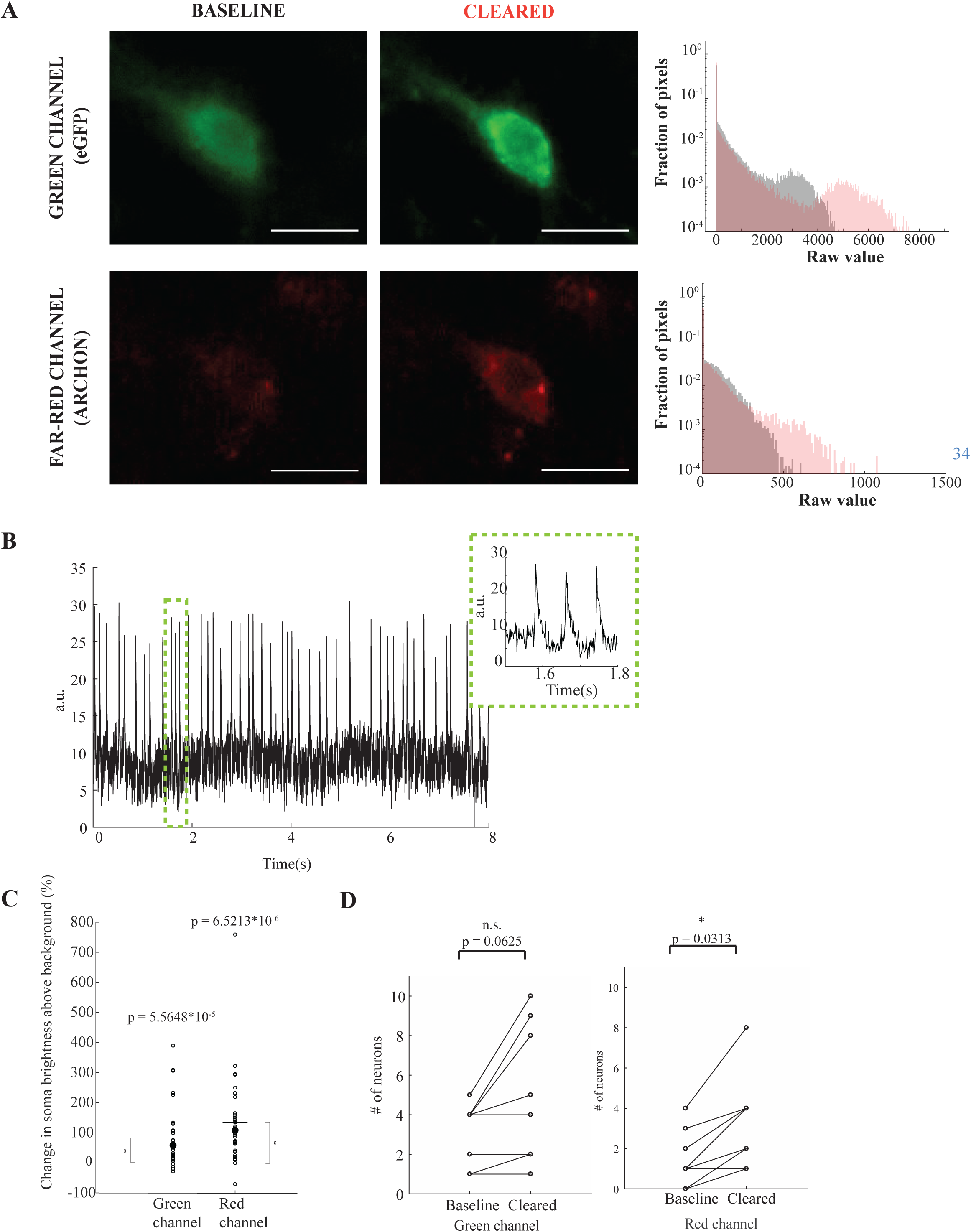
Voltage imaging in optically cleared acute cortical slices using the genetically encoded voltage indicator Archon-GFP. **A. Top:** a representative neuron imaged in the green channel (475/28nm excitation, emission 535/22nm band-pass filter) before (left) and after (right) optical clearing. **Bottom:** the same cell, imaged in the far-red channel (627nm excitation, emission 664nm long pass filter). Each pair of before and after images were acquired with the same settings and are shown using the same display settings. Scale bars = 10µm. The histogram for the raw pixel values is shown, plotted in semilogarithmic form, in the top right corner of each image. **B.** Representative trace of a spike train recorded from an optically cleared slice, while oxygenated iodixanol-aCSF was still being superfused. Enlargement of a representative 0.3s segment of the trace in the inset. **C.** Change in fluorescence intensity, after subtracting background, of cells expressing both GFP and Archon, imaged in the green and red channels before and after 1hr superfusion of the slices with iodixanol-aCSF with a refractive index 0.01 greater than plain aCSF. Reported p-values for the two-sided Wilcoxon signed rank test, n = 28 neurons from 8 slices, obtained from 4 mice. See **Results** for full statistics. **D. Left**: number of neurons visible in each slice before and after optical clearing, when imaging in the green channel Right: as on the left, for the far-red channel. In both sub-panels, reported p-values for the two-sided Wilcoxon signed rank test, 8 slices, obtained from 4 mice. See **Results** and **Supp. Table 9** for full statistics.

**Fig. 11.**
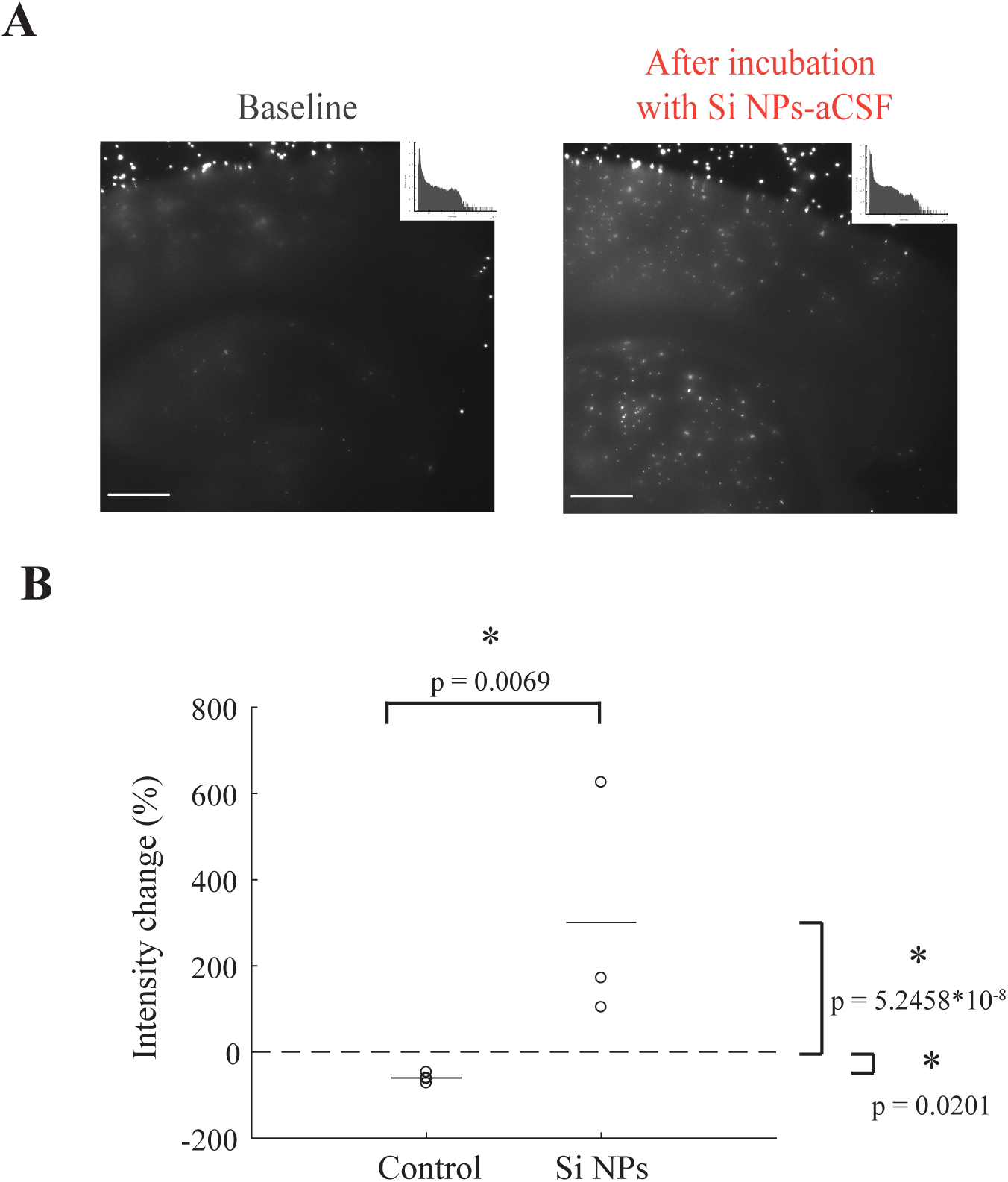
Initial testing of custom high-RI PEG-ylated silicon nanocrystals (Si NCs) for live tissue optical clearing. **A.** Array of 15μm diameter fluorescent (emission 645nm) polystyrene beads imaged though a 200μm-thick acute brain slice imaged with identical illumination and acquisition settings through a representative acute slice before (**left**) and after (**right**) incubation for 1hr in aCSF containing 50mg/ml Si NCs, with the same refractive index as the other solutions tested, under standard conditions for ex-vivo electrophysiology (see **Methods** for details). Display settings are the same for both conditions. Scale bar = 500µm. The histogram for the raw pixel values is shown, plotted in semilogarithmic form, in the top right corner of each image. **B.** Change in measured intensity of the signal from the beads imaged either through slices incubated in standard aCSF (control, also shown in Figure 1 **C., F., H**.) or Si NCs-aCSF. Each circle represents the average value for beads imaged through a slice. Solid line segments indicate means, the dashed line marks 0 (no change). See **Supp. Table 10** for full statistics, **Methods**, and **Discussion** for additional details and some considerations on the results and analysis.

Consistent with the results obtained when imaging neurons expressing the red fluorescent protein tdTomato, superfusion with dextran-aCSF significantly increased the signal that could be obtained from the neurons expressing the green genetically encoded calcium sensor compared to both baseline and superfusion with control aCSF. While we neither expected nor saw any evidence of increased overall levels of neural activity or of persistent intracellular depolarization resulting from application of the clearing agent in any of our *in vitro, ex vivo,* and *in vivo* studies (see **S.I. Figure 2**, **3**, and **4**), when imaging *in vivo* it is not possible, or at least extremely challenging, to fully measure the subthreshold dynamics and spiking patterns of every neuron in the volume of tissue imaged, so metrics assessing changes in measured signal from calcium sensors that could in principle be affected by changes in neural dynamics might be open to criticism. We therefore chose to focus on a metric that minimizes such theoretical risks, i.e. the lowest average somatic brightness above background observed over the course of a given recording, usually slightly over 10 minutes long (e.g. over the course of one set of visual stimuli presentation at baseline, or of one set of visual stimuli presentation post-superfusion, etc.), which can be thought of as the signal when the cell experiences minimal activity. This metric appeared consistent across different segments of a recording and across successive recordings for the same condition (e.g. for multiple visual stimulation sets recorded after superfusion, lasting overall several tens of minutes).

Since the effect of optical clearing observed in the previous experiments varied across depth (see **Figure 6** above), we only compared cells recorded at similar depths (in each experiment the imaging depth was chosen based on how active cells appeared to be in response to the presentation of visual stimuli), ranging approximately from 225 to 284µm (**Figure 9**; for dextran aCSF, median +183.2%, n = 73 cells from 3 mice; for control aCSF, median +30.4%, n = 81 cells from 2 mice; dextran aCSF vs control aCSF p = 1.8553*10^-45^, two-sample Kolmogorov-Smirnov test, see **Supp. Table 8** for full statistics). Small improvements seen after superfusion with control aCSF are likely attributable to the resolution of slight dural reddening at baseline observed in one control animal, which contributed approximately 58% of the control cells (47 out of 81).

### Functional imaging of a genetically encoded voltage indicator ex vivo following optical clearing

Genetically encoded voltage indicators allow the direct visualization of action potentials and subthreshold dynamics in vivo, but their practical use is constrained by the need to image with short exposures (300-1000 Hz imaging rates being typical) if full capture of high-speed dynamics is desired, and by the relative dimness of many of the sensors themselves. Live tissue optical clearing methods could therefore be beneficial for this application, since they could increase the magnitude of fluorescent signals from cells of interest within tissue. As a proof of this principle, we used iodixanol-aCSF, which had already shown promising results in the ex vivo bead assay (**Figure 1G**, **H** above) and in vivo (**Figure 3B, bottom panel** above), in conjunction with acute brain slices containing neurons that expressed the voltage sensor Archon-GFP^54^. Archon-GFP is a genetically encoded far-red voltage indicator fused with the green fluorescent protein GFP. As shown in **Figure 10A** for a representative neuron, after superfusion for one hour with iodixanol-aCSF, cells within the slices expressing Archon-GFP could be seen much more clearly, for the same imaging settings, in both the green (GFP) and red (Archon) channels, while the cells remained viable and could be successfully imaged (see **Figure 10B** for a representative trace).

All cells that could be visualized at baseline had approximate depth within 100-150µm from the surface of the slice, as expected from the use of a conventional epifluorescence microscope with an acute slice preparation. The signal above background in both the green and red channels increased significantly (**Figure 10C**; green channel, median +60.26%, p = 1.2569*10^-5^;; red channel, median +110.18%,, p = 1.8650*10^-7^,, N = 37 neurons from 11 sites from 9 slices from 4 mice, Wilcoxon signed-rank test; see **Supp. Table 9** for full statistical details). The number of neurons that could be seen in the slices also increased significantly, in the red channel, in which cells were less bright than in the green channel (**Figure 10D**; p = 0.0313 for the red channel, p = 0.0625 for the green channel, n = 37 neurons from 11 sites from 9 slices from 4 mice, Wilcoxon signed-rank test). Thus optical clearing may help with the visualization of voltage sensors and other cutting-edge reporters that may present unique challenges in imaging in intact brain circuitry.

### Ex vivo optical clearing employing high-RI silicon nanoparticles

Dextran, PEG, and iodixanol have relatively low refractive indices, on the order of ∼1.5, compared to ∼1.33 for extracellular fluid, so that millimolar concentrations of these reagents are required to raise the refractive index of the extracellular space appreciably. In order to achieve a greater degree of refractive index matching between the extracellular space and lipid membranes, while minimizing the concentration of material administered, the use of higher refractive index material could be advantageous. We here explored, at a proof-of-concept level, whether this might be possible.

Semiconductor nanoparticles, made from silicon and other elements, are promising candidates, because of the high refractive index of the native material (n = 4.5-5.6)^56^ and because of their limited toxicity compared to other nanomaterials^60,70^. Moreover, their surface can be functionalized in a variety of ways^71,72^, either to minimize interactions with biological molecules, or, potentially, to allow translocation across the cell membrane. Functionalizing the surface of nanoparticles with PEG has been used to enhance stability in vivo and biocompatibility of a variety of micro- and nanoparticles^73,73–75^. We synthesized PEG-ylated silicon nanocrystals (PEG-Si NCs), and found that at a concentration yielding a refractive index increase of ∼0.01 over plain aCSF, PEG-Si NCs could increase the transparency of acute brain slices (see **Figure 11**; median +171.94%, p = 5.2458*10^-8^, two-sample Kolmogorov-Smirnov test, n = 40, 69 beads for baseline and after optical clearing, respectively, from 3 slices from 2 mice, see **Supp. Table 11** for full statistical details). The changes observed for incubation with Si NCs-aCSF were significantly different from the changes observed with incubation in control aCSF (p = 0.0069, two-sample Kolmogorov-Smirnov test, respectively, n = 5 slices from 2 mice for control aCSF, n = 3 slices from 2 mice for Si NC-aCSF). Neurons in vitro that were exposed to the same concentration of PEG-Si NPs exhibited normal electrophysiological properties (**S.I. Figure 12**).

## Discussion

We here show that significant improvements in at-depth imaging can be achieved in the living brain through directly making the brain tissue itself more transparent, via refractive index mismatch reduction. Even though the refractive index mismatch between lipid membranes and aqueous compartments is thought to be on the order of 0.06-0.07, raising the refractive index of the artificial cerebrospinal fluid (aCSF) used to superfuse the cerebral cortex in vivo by only 0.01, through the addition of any of several different biocompatible materials, was sufficient to significantly enhance the signal above background from neurons in the mouse cortex, under both one-photon and two-photon imaging. The enhancement was especially appreciable deeper in the tissue where, at baseline and under control conditions, signals from cells were extremely low, or even indistinguishable from background. After clearing, many more cells became visible, and those that were identifiable at baseline showed, for the deepest range imaged (450 to 550µm), a median improvement in signal above background of ∼+385%, while many cells that were barely distinguishable at baseline showed increases of an order of magnitude or more. These results applied to all fluorophores tested, including green, red, and near-infrared fluorophores.

Key to our investigation was the safety of the procedure, given that many established methods for clearing fixed brain tissue or robust organs such as skin, bone, and ligament *in vivo* exist, but are incompatible with brain physiology. Comparing the visual tuning of neurons in the primary visual cortex of awake mice undergoing visual stimulation following superfusion with either control aCSF or dextran-aCSF showed no significant differences between the conditions, suggesting that basic functionality of complex neural networks in vivo might be preserved after optical clearing. These findings were reinforced by in vitro and ex vivo electrophysiological characterizations of neurons incubated in the same in vivo clearing agents, which showed no significant effect. We note that, as with any new technology, more extensive testing, with both positive and negative controls, would be helpful for gauging the utility of in vivo optical clearing in specific applications in everyday biology.

Going forward: although we chose to focus initially on cheap, widely available biocompatible materials, custom materials designed from the ground up for in vivo optical clearing could, in principle, offer a much greater degree of refractive index matching and, hence, efficacy, while causing minimal osmolarity increases. Silicon is an appealing material for this application because of its high refractive index (4.5, compared to 1.41-1.43 for dextran, PEG, and iodixanol), and low toxicity in vivo. We therefore developed PEG-surface functionalized silicon nanocrystals as a proof of principle demonstration, and showed their effectiveness ex vivo, obtaining an improvement, on average, of over 250% in the brightness above background of target beads imaged through an acute brain slice maintained under standard electrophysiology conditions. Beyond extracellular alteration, achieving full transparency may benefit from addressing intracellular refractive index mismatches, whether through exogenously delivered reagents, via genetically encoded methods (e.g., the potential adaptation, in the future, of refractive index modifying proteins for intracellular expression and refractive index matching^76^), or through hybrid methods.

In terms of practical applications: delivery of the clearing agent at depths greater than a few hundred microns is likely to require more complex approaches than the simple superfusion employed in this study; this problem has been extensively studied in the context of delivery of large molecules and nanoparticles for therapeutic applications, and a number of strategies, such as, for example, convection enhanced delivery (C.E.D.)^77,78^ or electrokinetic transport^79,80^ may be applicable to the delivery of optical clearing agents in vivo across large volumes of tissue.

Furthermore in this initial study, we focused on obtaining a conservative estimate of clearing agents’ effectiveness in vivo, when imaging static fluorophores during the experiments reported in this paper: we replaced the solution over the brain with standard aCSF for imaging; in practice, one could keep the cortex superfused with dextran-aCSF, to prevent any gradual washing out of the clearing agent from the tissue during the experiment. In addition, while the experiments here described were performed acutely, in many cases it would be advantageous to use a chronic preparation, with a head implant that allows superfusion of the cortex prior to/during imaging.

The recent adaptation of optogenetics for use in human therapeutics^81^ has generated considerable excitement about the possibilities of optically interfacing to the human brain. Live tissue optical clearing methods, in this regard, might also find more direct clinical use. One potential set of applications revolves around intraoperative optical histopathology and assessment of tumor margins (e.g. in conjunction with spectroscopic techniques^82–84)^, especially in tissues, such as the brain, where a conservative approach to tissue resection is particularly critical. Optical clearing methods designed to work without perturbing exquisitely delicate organs such as the brain may perhaps also be gentle enough to use in damaged tissues, for example to study and monitor the progression of healing in chronic wounds. It is at least encouraging that at least two of the reagents we tested, dextran and iodixanol, are in common clinical use for other applications, and easily available in pharmaceutical grade. While targeted safety assessments will have to be conducted, dextran, for example, appears to be safe, in an appropriate medium, when in direct contact with brain tissue through the cerebrospinal fluid, both acutely in non-human primates^85^ and chronically in rats^86^, at concentrations comparable to what would be used for optical clearing.

## ACKNOWLEDGMENTS

The Veinot Group thanks also thank the staff at Analytical and Instrumentation Laboratory in the Department of Chemistry at the University of Alberta for the assistance with FTIR analysis, and the University of Alberta Nanofab for support in material characterization.

## FUNDING

Lisa K. Yang (GTF, IG, ESB, DP, JPZ, NP, KP, DMA)

Howard Hughes Medical Institute (GTF, IG, ESB, DP, JPZ, NP, KP, DMA)

National Institutes of Health grant IR01MH123977 (GTF, IG, ESB, DP, JPZ, NP, KP, DMA)

National Institutes of Health grant R01DA029639 (GTF, IG, ESB, DP, JPZ, NP, KP, DMA)

National Institutes of Health grant R01MH122971 (GTF, IG, ESB, DP, JPZ, NP, KP, DMA)

National Institutes of Health grant RFINS113287 (GTF, IG, ESB, DP, JPZ, NP, KP, DMA)

National Institutes of Health grant 1R01DA045549 (GTF, IG, ESB, DP, JPZ, NP, KP, DMA)

John Doerr (GTF, IG, ESB, DP, JPZ, NP, KP, DMA)

Jeremy and Joyce Wertheimer (GTF, IG, ESB, DP, JPZ, NP, KP, DMA) Kavli Foundation (GTF, IG, ESB, DP, JPZ, NP, KP, DMA)

New York Stem Cell Foundation (GTF, IG, ESB, DP, JPZ, NP, KP, DMA)

U. S. Army Research Laboratory and the U. S. Army Research Office contract/grant number W911NF1510548 (GTF, IG, ESB, DP, JPZ, NP, KP, DMA)

NSF CBET grant 1344219

Friends of the McGovern Institute fellowship (GTF) Sheldon Razin Graduate Student Fellowship (IG)

National Institutes of Health grant ROIMH12635 (MH, MS) National Institutes of Health grant ROINS130361 (MH, MS) National Institutes of Health grant ROIMH133066 (MH, MS) National Institutes of Health grant K99EB027706 (MY)

National Institutes of Health grant R01DA045549 (HA, ZL, JG, UK)

Natural Science and Engineering Research Council (NSERC) Discovery Grant program (AI, PV, JV)

NSERC CREATE Alberta/Technical University of Munich International Graduate School for Hybrid Functional Materials (ATUMS, grant CREATE-463990-2015) (AI, PV, JV)

## AUTHOR CONTRIBUTIONS

Conceptualization: GTF, IG, ESB

Ex vivo and in vivo 1P imaging experiments: GTF, IG, with help from KP, YGY, NP

KP provided the viruses for the Archon-GFP experiments

2P in vivo imaging experiments: GTF, IG, MH, MY, with advice from MS In vitro and ex vivo safety assessments: IG, JPZ, NP

Initial theoretical analysis of refractive index changes: IG, AWG Denoising of the two-photon imaging data: ME, SH

Design and preparation of the Si nanocrystals: HA, ZL, JG, IG, U, AI, PV, JV Assessment of refractive changes in vitro: IG, ZQ, PS

Initial experiments in the project; IG, DMA

Analysis – discussion and design: GTF, IG, ESB, MS, YGY Analysis – design and implementation: GTF, with help from IG Visualization: GTF, with input and review by ESB

Funding acquisition: GTF, IG, DMA, ESB, MS, MY, UK, JV Supervision: ESB

Writing – original draft: GTF

Writing – review & editing: GTF, ESB

All authors contributed to the discussion of the data

## DATA AND MATERIALS AVAILABILITY

The data used in generating the figures in this paper as well as the images, static image stacks, and functional imaging datasets will be available upon request. The code for the SUPPORT denoising algorithm is available on https://github.com/NICALab/SUPPORT. All other custom code employed in the analysis will be available upon request.

