## Supplementary Information for "In Vivo Optical Clearing of Mammalian Brain"

Giovanni Talei Franzesi[1,2,3,4,5,6,7,8]†, Ishan Gupta[1,2,3,4,5,6,7,8]†, Ming Hu[4,9], Kiryl Piatkevich[1,3,5,10,11,12,13], Murat Yildirim[4,14], Jian-Ping Zhao[1,2,3,4,5,6,7,8], Minho Eom[15], Seungjae Han[15], Demian Park[1,2,3,4,5,6,7,8], Himashi Andaraarachchi[16], Zhaohan Li[16], Jesse Greenhagen[16], Amirul Muhammad Islam[17], Parth Vashishtha[17], Zahid Yaqoob[18], Nikita Pak[1,2,3], Alexander D Wissner-Gross[19], Daniel A. Martin-Alarcon[1,3,5], Jonathan J. Veinot[17], Peter T. C. So[3], Uwe Kortshagen[16], Young-Gyu Yoon[15], Mriganka Sur[4]‡, Edward S. Boyden[1,2,3,4,5,6,7,8]‡*

### Materials and Methods

All procedures involving animals were in accordance with the US National Institutes of Health Guide for the Care and Use of Laboratory Animals and approved by the Massachusetts Institute of Technology (MIT) Committee on Animal Care, , protocol numbers: 1221-099-24, 1218-100-21, 115-111-18, 0113-008-16, 1017-101-20.

Refractive index measurements

All refractive indices were measured with a Reichert 1310488M Abbe Refractometer (Cole-Parmer, IL).

Slice preparation for bead assays and ex vivo electrophysiology

C57BL/6 (Charles River Laboratory) or PV-cre mice (see below), either naïve or previously injected with a virus, were deeply anesthetized with a ketamine/xylazine cocktail, then transcardially perfused with ice-cold cutting solution containing, in mM, 252 sucrose, 3 KCl, 1.25 $NaH_2PO_4*2H_2O$, 2 $MgSO_4$, 2 $CaCl*2H_2O$, 10 glucose, 24 $NaHCO_3$, saturated with 95% oxygen and 5% carbon dioxide. The brain was then extracted and slices 200, 250, or 300 µm thick were cut on a vibratome (Leica Biosystems). The slices were then allowed to recover for at least 30' in an interface holding chamber (Harvard Apparatus), in recording aCSF, containing, in mM, 126 NaCl, 3 KCl, 1.25 $NaH_2PO_4*2H_2O$, 2 $MgSO_4$, 2 $CaCl*2H_2O$, 10 glucose, 24 $NaHCO_3$, and saturated with 95% oxygen and 5% carbon dioxide**.** The brain was then extracted and slices 200 (for Si nanoparticle testing), 250 (for PEG testing), or 300 (for electrophysiology) µm thick were cut on a vibratome (Leica Biosystems, Wetzlar, Germany). The slices were then allowed to recover for at least 30' in an interface holding chamber (Harvard Apparatus), in aCSF containing, in mM, 126 NaCl, 3 KCl, 1.25 $NaH_2PO_4*2H_2O$, 2 $MgSO_4$, 2 $CaCl*2H_2O$, 10 glucose, and 24 $NaHCO_3$, and saturated with 95% oxygen and 5% carbon dioxide. Optical clearing solutions were similar to recording aCSF, with the addition of either 40mg/ml Dextran 40kDa, 60mg/ml PEG10kDa, 40mM Iodixanol, or a sufficient concentration of Si NPs to obtain a refractive index increment of ~0.01. All chemicals, other than the Si NPs, were obtained from Millipore Sigma (MO)

Bead assay

Acute brain slices, prepared as described above, were placed in a sealed imaging chamber (VWR) filled with plain aCSF. An array of red (emission, 645 nm) fluorescent 15 µm polystyrene beads (Invitrogen, MA) was then imaged through the slice on an inverted epifluorescence microscope. The slices were then transferred to an incubation chamber containing the clearing solution (e.g. Dextran-containing aCSF), which was constantly bubbled with 95% oxygen, 5% carbon dioxide. After one hour the slices were removed from the incubation chamber, rinsed once with plain aCSF, and placed in the sealed imaging chamber, filled with plain aCSF. We then imaged the beads through the slice again, using the same exposure and light intensity settings as before clearing. Data was analyzed using Fiji(*69–71*). See below for details on the data analysis.

In vitro whole-cell patch clamping

Whole-cell patch-clamp recordings were made using an Axopatch 200B amplifier, a Digidata 1440 digitizer, and a PC running pClamp10.4 (Molecular Devices, CA). For current clamp recordings, neurons were patched at 18-21 days in vitro (DIV) to allow for sodium channel maturation. Neurons were bathed in room-temperature Tyrode's solution containing, in mM, 129 NaCl, 2 KCl, 3 CaCl2, 1 MgCl2, 25 HEPES, 30 glucose, 0.01 NBQX, 0.01 GABAzine and pH adjusted to 7.3 with NaOH or in Tyrode's solution supplemented with 40mg/ml Dextran 40kDa, 60mg/ml PEG 10kDa (Dextran-Tyrode and PEG-Tyrode, respectively), or with 40mM Iodixanol (Iodixanol-Tyrode). Neurons were bathed in Tyrode's solution or modified Tyrode for 1 hour or two hours prior to whole-cell patch clamping (for the one hour and two hours data sets, respectively). Borosilicate glass pipettes (Warner Instruments, CT) with an outer diameter of 1.2 mm and a wall thickness of 0.255 mm were pulled to a resistance of 5–9 MΩ with a P-97 Flaming/Brown micropipette puller (Sutter Instruments) and filled with a solution containing, in mM, 150 K-gluconate, 8 NaCl, 0.1 CaCl2, 0.6 MgCl2, 1 EGTA, 10 HEPES, 4 Mg-ATP, 0.4 Na-GTP, with pH adjusted to 7.3 with KOH, and osmolarity adjusted to 298 mOsm with sucrose. We used cells with access resistance 5-30 MΩ and holding current within ±50 pA (at −65 mV, in voltage clamp). Access resistance was monitored throughout recording. Data were analyzed using Clampfit (Molecular Devices, CA) and custom Matlab scripts (MathWorks, MA).

Primary neuronal culture

Hippocampal neurons were prepared from postnatal day 0 or 1 Swiss Webster (Taconic) mice as previously described(*72*) but with the following modifications: dissected hippocampal tissue was digested with 50 units of papain (Worthington Biochem, NJ) for 8 min, and the digestion was stopped with ovomucoid trypsin inhibitor (Worthington Biochem, NJ). Cells were plated at a density of 16,000–40,000 per glass coverslip coated with Matrigel (BD Biosciences). Neurons were seeded in 90 or 100ul Plating Medium containing MEM(Life Technologies), glucose (33mM, Sigma), transferrin(0.01%, Sigma), Hepes (10mM, Sigma), Glutagro(2mM, Corning), Insulin (0.13%, Millipore), B27 supplement (2%, Gibco), heat inactivated fetal bovine serum (7.5%, Corning). After cell adhesion, additional Plating Medium was added. AraC (0.002mM, Sigma) was added when glia density was 50-70%. Neurons were grown at 37C degree and 5% CO2 in a humidified atmosphere.

Ex vivo whole-cell patch clamping

Slices were prepared as described above (see *Slice preparation for bead assays and ex vivo electrophysiology).* The slices were constantly superfused with carbogenated aCSF for baseline recordings, then with carbogenated dextran-aCSF (for two hours and for the subsequent recordings). Patch-clamp recordings under DIC guidance were performed as described above for in vitro whole cell patch clamping (see *In vitro whole-cell patch clamping*).

Ex vivo imaging of the genetically encoded voltage sensor Archon-GFP

Slices were prepared as described above (see *Slice preparation for bead assays and ex vivo electrophysiology).* We first imaged areas within the slices while superfusing them with regular artificial cerebrospinal fluid (aCSF) saturated with 95% oxygen, 5% carbon dioxide. We then superfused them for one hour with aCSF containing 6% iodixanol (Iodixanol-aCSF), also saturated with 95% oxygen, 5% carbon dioxide, and imaged the same regions again. Finally, we imaged the voltage dynamics of individual neurons, while still superfusing the slices with Iodixanol-aCSF. Neurons were imaged in the green (475/28 nm excitation, emission 535/22 band pass filter) channel before (left) and after (right) optical clearing, and the same was done for the far red (excitation 637nm, emission 664 long pass filter) channel before and after optical clearing. The images were taken with the same illumination and exposure. The equipment used consisted of a

Nikon Eclipse Ti inverted microscope used in conjunction with a 40 × NA 1.15 water immersion objective (Nikon, Japan), a 637-nm Laser (637 LX, OBIS, NH) focused on the back focal plane of the objective, a SPECTRA X light engine (Lumencor, OR) having 475/28 nm, 585/29 nm, and 631/28 nm exciters (Semrock, IL), a 470 nm LED (ThorLabs, NJ) and a 5.5 Zyla camera (Andor, MA), controlled by NIS-Elements AR software.

Transgenic animals

*Mice expressing tdTomato in PV+ neurons.* Transgenic animals expressing the Cre recombinase under control of the parvalbumin promoter (B6.129P2-Pvalbtm1(cre)Arbr/J, commercially available from The Jackson Laboratory, ME, abbreviated as PV-cre in the rest of the section) were crossed with Ai14 mice (B6.Cg-Gt(ROSA)26Sortm14(CAG-tdTomato)Hze/J, commercially available from The Jackson Laboratory, ME), in order to express the tdTomato fluorophore in parvalbumin-positive neurons.

*Mice expressing calcium sensors in excitatory nerons.* Transgenic animals expressing genetically encoded fluorescent calcium sensors in excitatory neurons were generated either by crossing mice expressing the Cre recombinase under the control of the CamkII promoter with Ai148 and Ai148d(for GCaMP6f) lines, or by crossing Emx-1-IRES-cre animals with Ai93 (for GCaMP6f) or Ai94 (for GCaMP6s) lines. All transgenic lines were obtained from The Jackson Laboratory, ME. All mice used in the experiments described were adults (> 8 weeks old), and both male and female mice were used.

Virus injection

In order to express iRFP682(*73*) or the genetically encoded voltage indicator Archon-GFP(*36*) in a sparse subset of neurons we injected a mixture of dilute (1:500) AAV2/8-CAG-Cre and AAV2/8-FLEX-iRFP682 or AAV2/8-FLEX-Archon-GFP, respectively, in C57BL/6 mice, AAV2/8-FLEX-iRFP682 alone in PV-cre animals, or AAV2/8-FLEX-Archon-GFP in PV-cre animals. Under sterile conditions, a small craniotomy was performed, and 0.5-1.5µl of the viruses were injected at a site -1.5 mm posterior (A/P) to bregma and 1.5 mm lateral (M/L) from bregma at a depth of 0.15-0.2 mm. We then allowed at least 4 weeks for the protein to be expressed. For expressing the genetically encoded voltage indicator Archon-GFP, a similar procedure was followed.

In vivo one photon imaging

On the day of the experiment, with the animal under isoflurane anesthesia (1-2% in $O_2$), a metal headplate was secured to the skull using dental cement. A small (<1mm) craniotomy was then performed at the target site in S1 or V1 (approximately -1.5 A/P, 1.5 M/L and -3mm A/P, 2.5mm M/L from bregma, respectively), followed by dural permeabilization via agarose beads functionalized with collagenase(*74*). The animal, still kept under isoflurane anesthesia, was then transferred to the custom-built one-photon microscope. For tdTomato imaging we used 531nm excitation with a 580nm long pass emission filter (Thorlabs, NJ), while for iRFP imaging we used 625nm excitation, with a 664 long pass emission filter (Thorlabs, NJ). The headplate holder was secured on a platform mounted on a 3-axis stage, which could be controlled through Micromanager to acquire image stacks. After a baseline stack was taken, with the brain covered with lactated Ringer's solution, the solution of interest (saline, sucrose -aCSF, Dextran- aCSF, PEG-aCSF, or Iodixanol-aCSF) was superfused onto the brain. Every 20 minutes, for one hour, it was removed and replaced with fresh solution, to minimize the effects of evaporation. At the end of the hour the solution was removed, the surface rinsed once with lactated Ringer's solution, then fresh lactated Ringer's solution was applied and a new image stack was taken. The composition of the HEPES-aCSF was, in mM: 135 NaCl, 5 KCl, 5 HEPES, 1.8 $CaCl_2*2H_2O$, 1 $MgCl_2*6H_2O$. Dextran-HEPES-aCSF contained, in addition, 40mg/ml Dextran 40kDA. PEG-HEPES_aCSF contained, in addition, 60 mg/ml PEG10kDa, whereas sucrose-HEPES-aCSF contained, instead of PEG10KDa, 6 mM sucrose, as an osmotically matched control.

In vivo two-photon imaging

For two-photon in vivo imaging the surgical preparation of the animal was the same as for one photon imaging (see above). For static imaging of tdTomato labeled cortical cells the animal was transferred to a custom headplate holder mounted under the microscope objective while still under anesthesia, which was maintained with 0.5-1% isoflurane in $O_2$ for the duration of the experiment. For awake functional imaging, after the animal was transferred and secured to the imaging headplate holder, it was allowed to recover for 20-30' before the acquisition of the baseline visual stimulation recordings commenced. In both kinds of experiments imaging, at baseline as well as after superfusion, was done under standard aCSF. In control animals, the cortex was kept continuously superfused with control aCSF, while for clearing experiments, after acquisition of the baseline images the cortex was superfused with dextran-aCSF, which was removed and replaced every 25-35' in order to minimize any possible change in osmolarity and dextran concentration due to evaporation. Prior to post-clearing imaging the cortex was rinsed twice with standard aCSF, before fresh standard aCSF was applied.
Imaging was carried out with a custom two-photon laser-scanning microscope (Ti:Sapphire, Mai-Tai, Spectra Physics; modified Fluoview confocal scan head, ×20 lens, 0.95 NA, Olympus). Excitation for fluorescence imaging employed 100-fs laser pulses (80 MHz) at 920 nm for GCaMP-family calcium sensors and 1,020 nm for tdTomato. Images were acquired in resonant mode, averaging for images per frame, at an effective imaging rate of ~7.5 Hz. A 2x digital zoom was employed.

Visual stimuli presentation

Visual stimuli were delivered via a 17 inch LCD display placed 15 cm from the eyes. Stimuli were generated in Matlab (Mathworks, Natick, MA), using the PsychoPhysics Toolbox (*4*). Square wave drifting gratings with 100% contrast were used to test orientation tuning. Grating stimuli were presented in pseudorandom order, with each 3s stimulus followed by 2s blank gray screen presentation. Each stimulus was presented 10 times for each recording.

Ex vivo bead assay data analysis

We compared the intensity of the signal from the beads that could be seen through the slice at baseline to that of the beads that could be seen through the same slice after optical clearing or incubation with regular aCSF. Since the ex vivo bead assay was meant as a screening step to identify promising candidate for in vivo optical clearing, we focused on M1, SI, V1 cortical regions. For each slice, we first adjusted brightness and contrast to maximize visibility of the beads through the slice at baseline, and drew a circular ROI in Fiji around each visible bead within the contours of the slice. We then went back to the raw image and calculated the average intensity of the signal in each bead ROI. When analyzing the images for the same slices after incubation in the clearing agent we proceeded in the same way, but restricted our analysis to beads within those areas of the slice were beads were also visible at baseline, to ensure that the two sets of measurements were comparable (after clearing, beads were often visible in other regions, too, but we lacked a baseline measurement to compare them with). Even using far-red beads to maximize their visibility through slices thick enough to remain viable reliably, the intrinsic opacity of the tissue, variable from location to location within the slice, especially at baseline and after incubation in control aCSF, only allowed some beads to be seen.

Denoising of two-photon static imaging data

The imaging stacks acquired above were denoised using SUPPORT(*75*), which removed the Poisson–Gaussian noise in the images by learning and utilizing the spatial and temporal dependence among the pixel values (publicly available on https://github.com/NICALab/SUPPORT), to denoise baseline and post-superfusion images. For processing two-photon static imaging data, we employed a single network for training and testing on all z-stack images. The training process involved 100 epochs with a batch size of 64 and utilized patches of size 5(z) × 128(x) × 128(y). The SUPPORT network had a receptive field size of 5(z) × 148(x) × 148(y) and a blind spot size of 1(z) x 3(x) × 3(y). For processing two-photon functional imaging data, we utilized separate networks for training and testing on each recording. These networks were trained for 10 epochs with a batch size of 64, using patches of size 5(t) × 128(x) × 128(y). The SUPPORT network employed a receptive field size of 5(t) × 168(x) × 148(y) and a blind spot size of 1(t) × 23(x) × 3(y). All computations were performed on a workstation equipped with two Intel Xeon Gold 6226R CPUs, 128 GiB of RAM, and an NVIDIA GeForce RTX 4090 GPU.

In vivo imaging data analysis

The analysis proceeded in two main steps: identifying the cell bodies of cells visible in both conditions and selecting the corresponding regions of interest (ROIs), and then calculating the difference in mean brightness between the cell body and its immediate neighborhood in the raw images. The former was performed using Fiji(*69–71*), the latter using custom-written MATLAB code.

To facilitate identifying the cell bodies, we first subtracted the background (as described below) for each image in the stack, then summed overlapping sets of 5-20 images. We first smoothed each image with a Gaussian blur, with a radius of that was much larger than the size of the cell bodies, yet not so large that it obliterated differences in background brightness across the imaging plane. Empirically, some ranges of values seemed to yield images that were more useful for the subsequent analysis. While values ~50% smaller or ~100% larger also worked well, we settled on values of 200,100, or 40 pixels for images that had dimensions of 2048x2048 (two-photon imaging, 2P), 1024x1024 (one-photon imaging, 1P), or 512x512 pixels (one-photon imaging, 1P), respectively. The resulting background image was then subtracted from each original image in the stack. For two-photon imaging stacks, the resulting, background-subtracted image was further smoothed with a Gaussian filter with a very small (~5% of cell body diameter) radius. Overlapping sets of 5-20 images were then summed, using the z-project function Fiji. The somatic locations were then manually selected using elliptical selection tool in Fiji, either on the summed images or in individual background subtracted slice sin the stack, since using both approaches allowed more reliable identification of cell bodies, especially when cells were dime and/or very close to each other in XY coordinates. Using the “fit ellipse” function in Fiji, ellipses were then fit to the regions of interest (ROIs) and the results exported as excel files. The analysis was done with the (blinded) image stacks for two conditions side by side, in order to ensure that cell identification was consistent between them for paired analysis. For a subset of the cells, the ROI identification was performed twice, several months apart, as well as for each image stack separately, with cell identification being matched in a separate, subsequent step, to check for robustness.

After having identified the ROIs corresponding to the cell bodies in this way, we went back to the original images (raw images for 1P data, SUPPORT-denoised, as described above, images for 2P data) to calculate how bright the cell bodies were compared to the area immediately around them. This was a better metric than simply the brightness of the cell bodies, especially under 1P imaging, because, due to scattering, autofluorescence, local variations in density of labeled cells, etc., different areas of the image could have very different levels of background brightness, so that, for example, a cell body of a given brightness could stand out sharply in one region, but be indistinguishable from background in another.

The analysis was performed using custom-written MATLAB code. For each ROI, the code calculated the difference in intensity values between the soma and the area around it over an interval of 10-20 slices from its putative z location within the z-stack, and returned the highest value for this difference, as well as the image stack index (depth) this corresponded to. If there was a significant mismatch in the depth value corresponding to the maximum soma vs background intensity difference for the image stacks corresponding to the before and after incubation conditions, after controlling for brain deformation and changes in overall starting points common to all cells within an image, the cell was discarded and not included in the analysis, since the result was likely due to very bright neighboring cells immediately above or below the actual location of the cell body of interest. This only happened in a few cases (<20 cells across all animals). Since during imaging there could be some degree of brain motion, and the XY coordinates of a given cell body could therefore shift slightly across the z stack, we excluded from the background are a thin (~10-20% of the soma diameter) ring immediately around the identified cell body. Small changes in the ring width or width of the background area didn't affect the results (<1% change).

<u>Inclusion/exclusion criteria, attrition, blinding, and randomization</u>
Animals were excluded from the analysis if there were surgical or post-surgical issues (e.g. bleeding) that could confound the results. A small number of animals (3) didn't have any visible fluorescently labeled cells at baseline, because of variable expression levels, and could not be used for the imaging and superfusion experiments. The analysis was performed with the experimenter blind to the category (control vs cleared and baseline vs. post-superfusion/post-incubation) of the image set. For a given type of image stack (e.g. acquired under one photon microscopy or acquired on a different scope under two photon microscopy) the same routine for extracting the cell body and background values (see Methods) was run for all stacks, regardless of condition (control vs. cleared).

<u>Statistical tests</u>
Since the data generally was not, and could not be assumed to be, normally distributed, we employed the following well-established non-parametric tests: the Wilcoxon signed-rank test, for

before/after comparisons, and the two-sample Kolmogorov-Smirnov test for comparing independent distributions.

Visual tuning analysis

The ROIs of the cells were identified as described above (see *In vivo imaging data analysis*), and the average intensity of each ROI in each frame was extracted in Fiji. After aligning the stimulation data and the recorded images using custom-written Matlab code (MathWorks, IL), The section of recording corresponding to each stimulus was identified. The lowest recorded value over time for average intensity in each ROI was used as baseline fluorescence for calculating the $\Delta F/F_0$ response to each type of visual stimulus. Orientation Selectivity Index and Direction Selectivity Index values were calculated as in(*50*, *51*).

Silicon nanocrystals (Si NCs) synthesis and functionalization

*Si nanocrystal (Si NC) synthesis*: Si NCs were synthesized in a continuous-flow, low pressure, nonthermal plasma reactor from an argon-silane (Ar-$SiH_4$) gas mixture using a previously published procedure(*76*), 30 standard cubic centimeters (sccm) of Ar and 14 sccm of 5% $SiH_4$ diluted in helium were introduced through ¼” borosilicate glass reactor tube, which expanded to a diameter of 1” with an inlet for 100 sccm of $H_2$ injection. The subsequent $H_2$ injection was used to facilitate surface passivation of synthesized Si nanocrystals. The plasma was generated in the reactor by the application of nominal (read at the power source) 50 W radiofrequency power coupled through an impedance matching network to two copper electrodes positioned 4 cm above the $H_2$ injection point. The frequency was 13.56 MHz. The gas pressure was maintained at ~1.5 Torr through an adjustable downstream orifice, which also served to accelerate the nanocrystals for collection by impaction on glass substrates.

*Si NC surface functionalization:* As synthesized Si NCs (H-Si) were subsequently subjected to a solution phase reaction with polyethylene glycol methyl ether acrylate ligands (PEG) (Sigma Aldrich #454990) to form water dispersible Si NCs. All solvents used for surface functionalization were dried on molecular sieves (Millipore Sigma #208604, type 4A) and were degassed through nitrogen bubbling, unless otherwise noted. First, as synthesized Si NCs were transferred to a nitrogen filled glovebox via a load-lock system to avoid surface oxidation. Fresh Si NC powder was mixed with a mixture of mesitylene (Sigma Aldrich, M7200, 98%) and PEG ligand to form a suspension. For one set of experiments, roughly 5 mg of Si NCs were mixed with 4.85 mL of mesitylene and 0.15 mL of PEG ligand in a pressure vial and heated at 160 °C for 16 h inside a nitrogen filled glove box. The products were isolated and excess ligands were removed by rinsing with hexane (Sigma Aldrich #208752, > 95%) followed by three times centrifugation. Isolated PEG grafted Si NCs were dried under vacuum and used for clearing experiments.

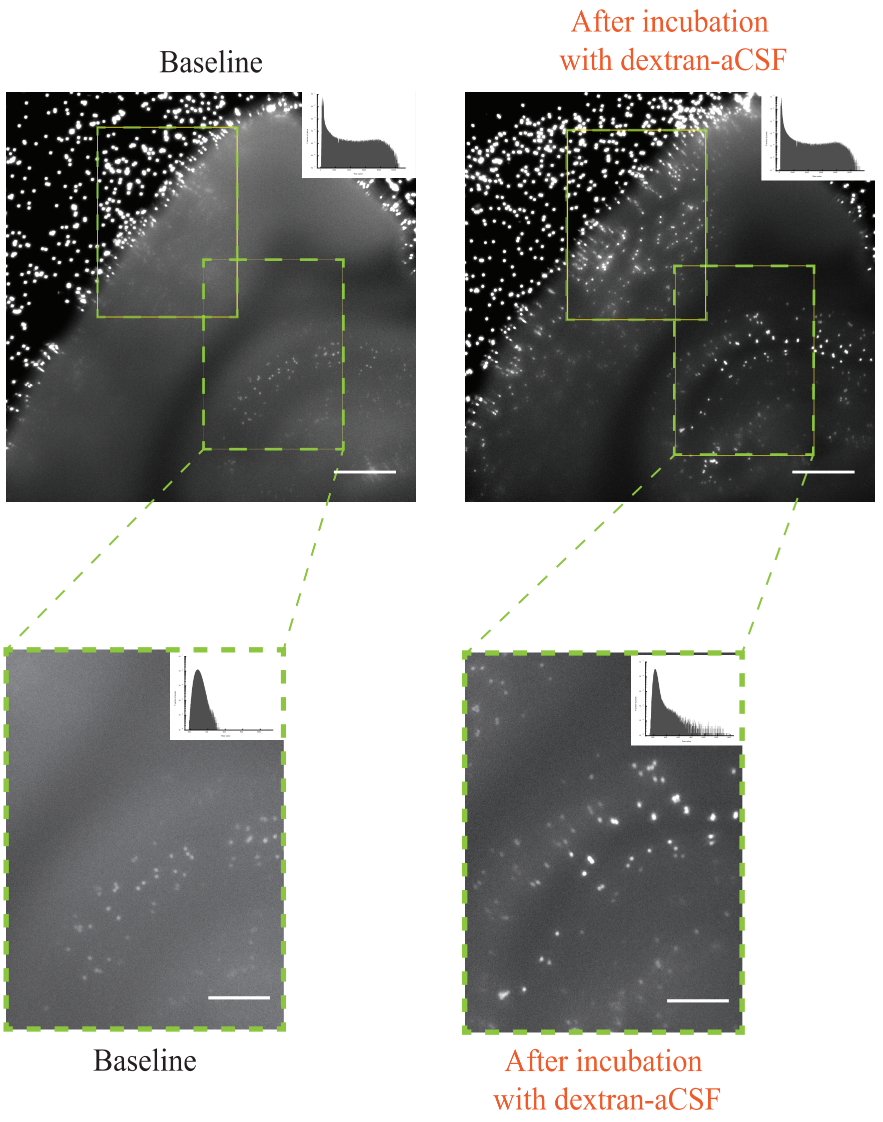


**Fig. S1.**

**Example of optical clearing outside of cortex (hippocampus) with dextran-aCSF. A.** Array of fluorescent (emission 645nm) 15µm polystyrene beads imaged though a 250µm-thick acute brain slice imaged with identical illumination and acquisition parameters before (**left**) and after (**right**) incubation for 1hr in dextran-containing aCSF under standard conditions for *ex vivo* electrophysiology (see **Methods** for details). Display settings are the same for both conditions. The histogram for the raw pixel values is shown, plotted in semilogarithmic form, in the top right corner of each image.

**A**

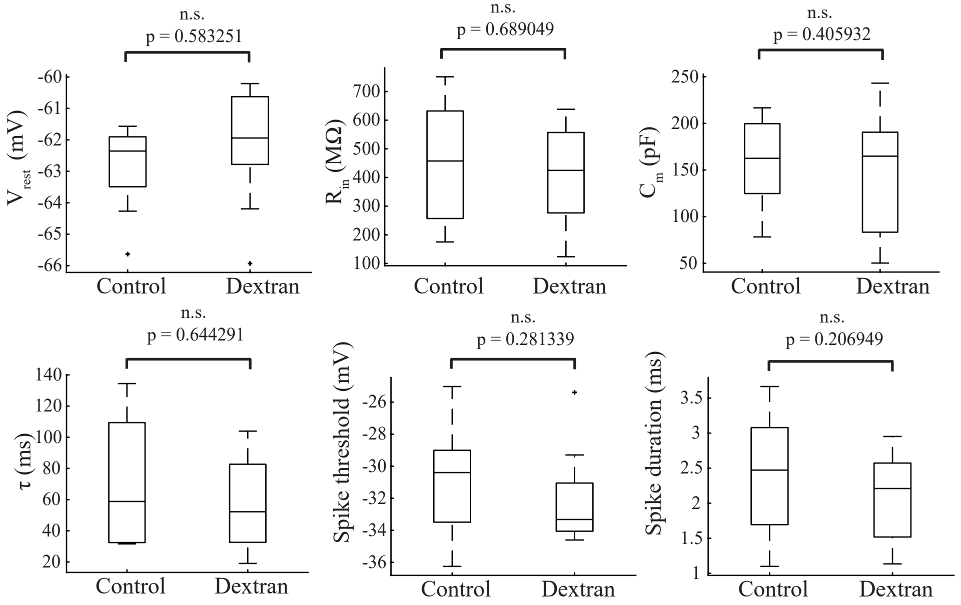
n.s.
p = 0.583251
$V_{rest}$ (mV)
Control
Dextran
n.s.
p = 0.689049
$R_{in}$ (MΩ)
Control
Dextran
n.s.
p = 0.405932
$C_m$ (pF)
Control
Dextran
n.s.
p = 0.644291
τ (ms)
Control
Dextran
n.s.
p = 0.281339
Spike threshold (mV)
Control
Dextran
n.s.
p = 0.206949
Spike duration (ms)
Control
Dextran

**B**

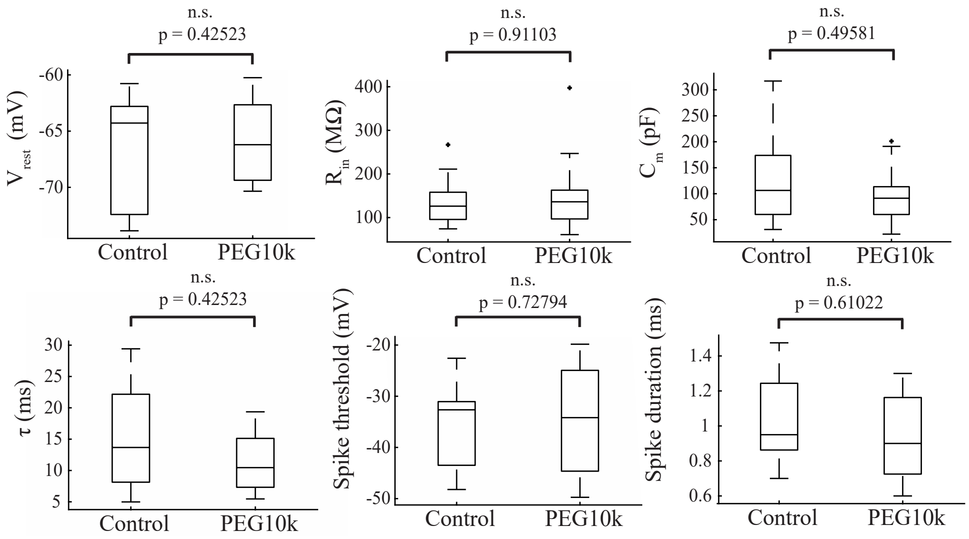
n.s.
p = 0.42523
$V_{rest}$ (mV)
Control
PEG10k
n.s.
p = 0.91103
$R_{in}$ (MΩ)
Control
PEG10k
n.s.
p = 0.49581
$C_m$ (pF)
Control
PEG10k
n.s.
p = 0.42523
τ (ms)
Control
PEG10k
n.s.
p = 0.72794
Spike threshold (mV)
Control
PEG10k
n.s.
p = 0.61022
Spike duration (ms)
Control
PEG10k

**C**

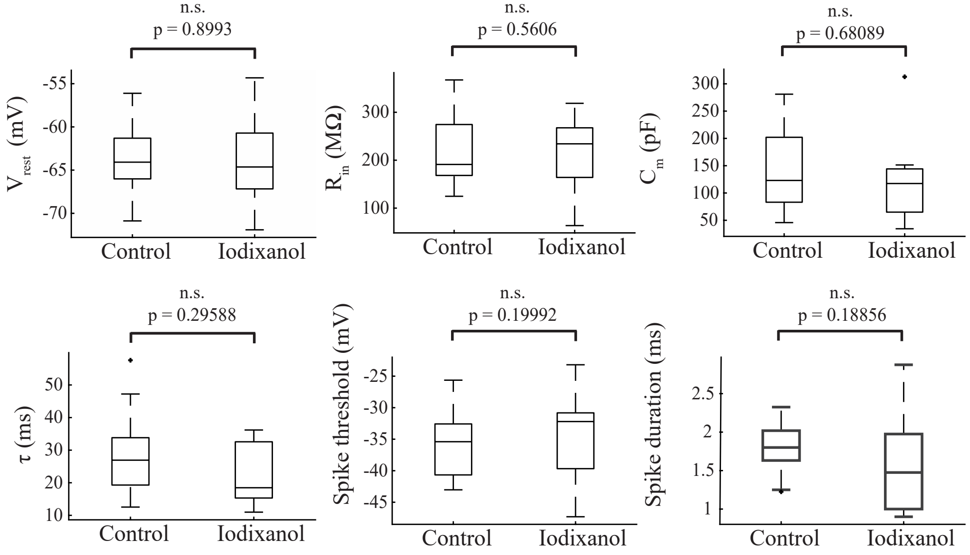
n.s.
p = 0.8993
$V_{rest}$ (mV)
Control
Iodixanol
n.s.
p = 0.5606
$R_{in}$ (MΩ)
Control
Iodixanol
n.s.
p = 0.68089
$C_m$ (pF)
Control
Iodixanol
n.s.
p = 0.29588
τ (ms)
Control
Iodixanol
n.s.
p = 0.19992
Spike threshold (mV)
Control
Iodixanol
n.s.
p = 0.18856
Spike duration (ms)
Control
Iodixanol

**Fig. S2.**

**Electrophysiological properties of primary neuronal cultures are preserved after one hour incubation with Tyrode containing the same concentrations and reagents used for live tissue optical clearing ex vivo and in vivo. A.** Comparison of electrophysiological properties assessed by patch clamping of neurons in culture, incubated for one hour either in standard Tyrode or in Tyrode containing 1.5mM Dextran 40kDa. From top left: resting membrane potential, input resistance, membrane capacitance, membrane time constant, spike threshold, and spike duration. Control: n = 12 neurons from 5 cultures. Dextran: n = 11 neurons from the same 5 cultures as for control. **B.** as in **A.**, for Tyrode containing 6mM PEG 10kDa. n = 15 cells from one culture for treated cells, n = 17 cells for control cells, from the same culture. **C.** As in **A.** and **B.**, for Tyrode containing 40mM iodixanol. n = 15 neurons from 3 cultures for control, n = 14 neurons from 2 cultures for iodixanol-Tyrode. Throughout the figure, n.s. = not significant, at the 5% level. All reported p-values calculated using the two-sample Kolmogorov-Smirnov test.

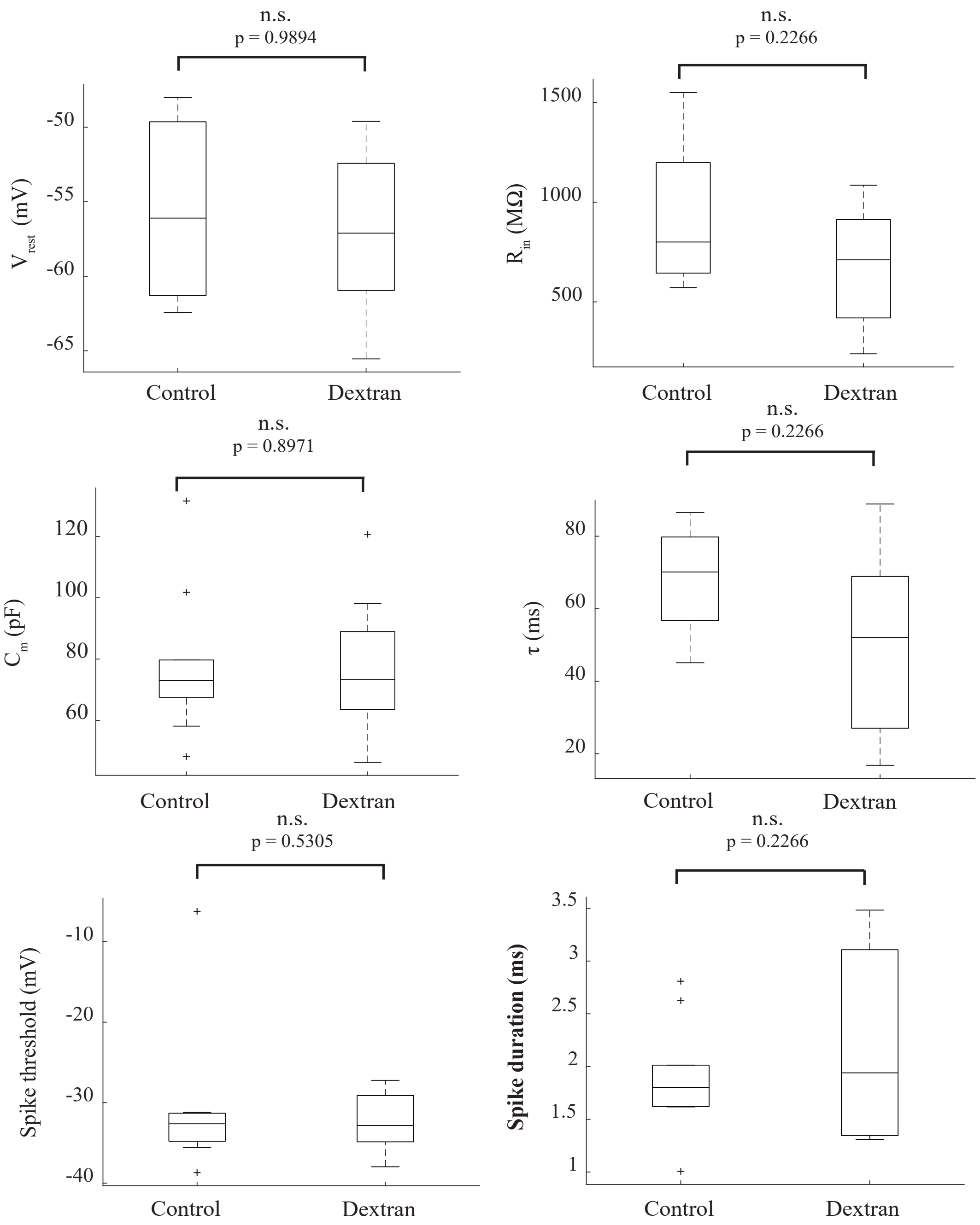


**Fig. S3.**

**Electrophysiological properties of primary neuronal cultures are preserved after two hours incubation with Tyrode containing the same concentration of Dextran 40kDa as used for in vivo and ex vivo optical clearing.** Comparison of electrophysiological properties assessed by patch clamping of neurons in culture, incubated for two hours either in standard Tyrode or in Tyrode containing 1.5mM Dextran 40kDa. From top left: resting membrane potential, input resistance, membrane capacitance, membrane time constant, spike threshold, and spike duration. Control: n = 19 neurons from 3 cultures. Dextran: n = 12 neurons from the same 3 cultures as for control. Throughout the figure, n.s. = not significant, at the 5% level. All reported p-values calculated using the two-sample Kolmogorov-Smirnov test.

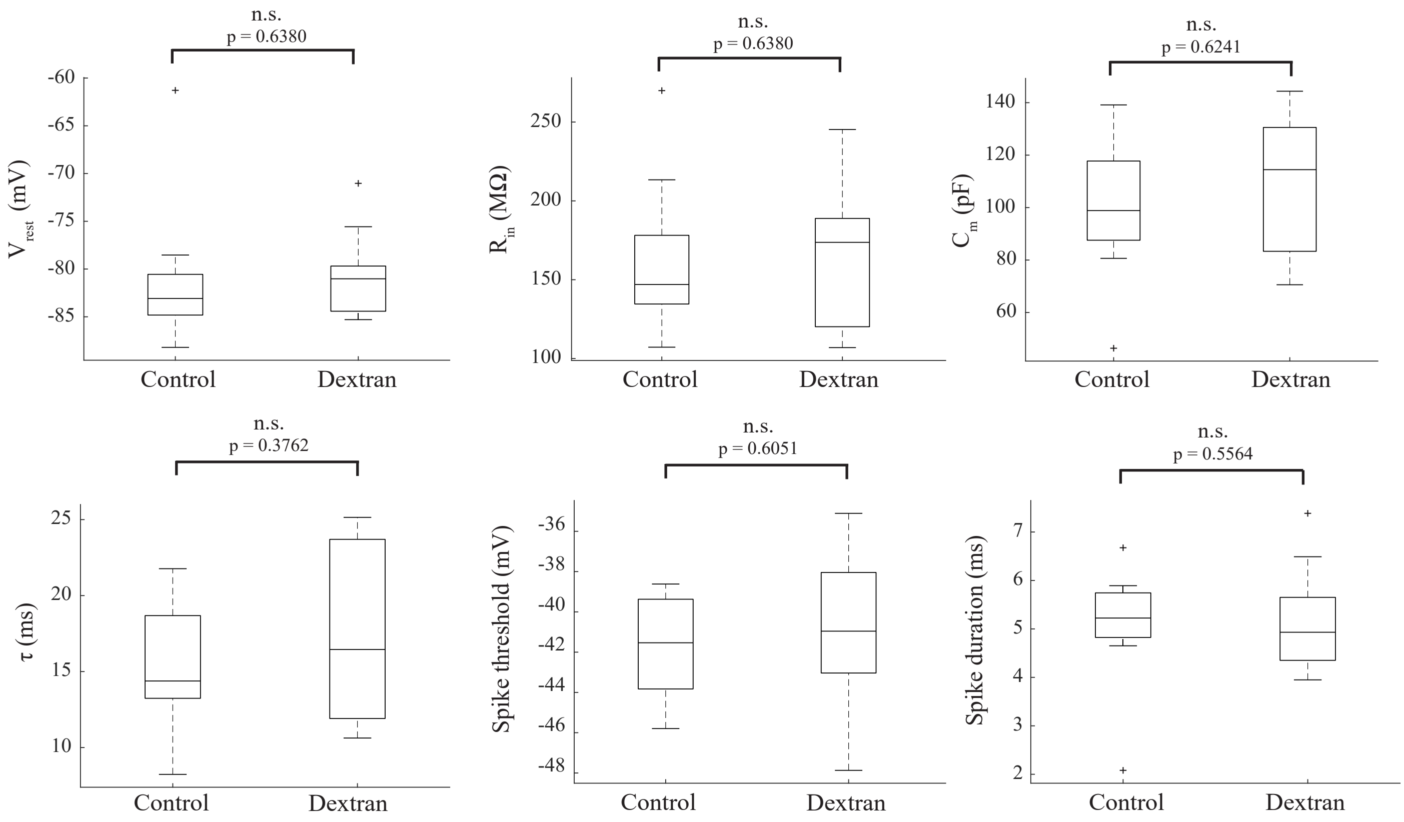


**Fig. S4.**

**Electrophysiological properties of cortical neurons in acute slices are preserved after two hours superfusion with Dextran-aCSF.** Comparison of electrophysiological properties assessed by patch clamping of hippocampal pyramidal cells in acute slices either before or after two hours superfusion with aCSF containing 1.5mM Dextran 40kDa. From top left: resting membrane potential, input resistance, membrane capacitance, membrane time constant, spike threshold, and spike duration. For the control/baseline condition, n = 12 cells; for the after superfusion with dextran-aCSF condition, Dextran: n = 11 cells, in both cases from 5 slices obtained from 5 animals. Throughout the figure, n.s. = not significant, at the 5% level. All reported p-values calculated using the two-sample Kolmogorov-Smirnov test.

**A**

### CONTROL ACSF

*Display settings autoscaled to the post-superfusion image of each pair and applied to the paired baseline and post-superfusion images.*
*Note that each ROI I-III was autoscaled separately based on the post-superfusion ROI image, with the same settinsg then applied to the baseline image*

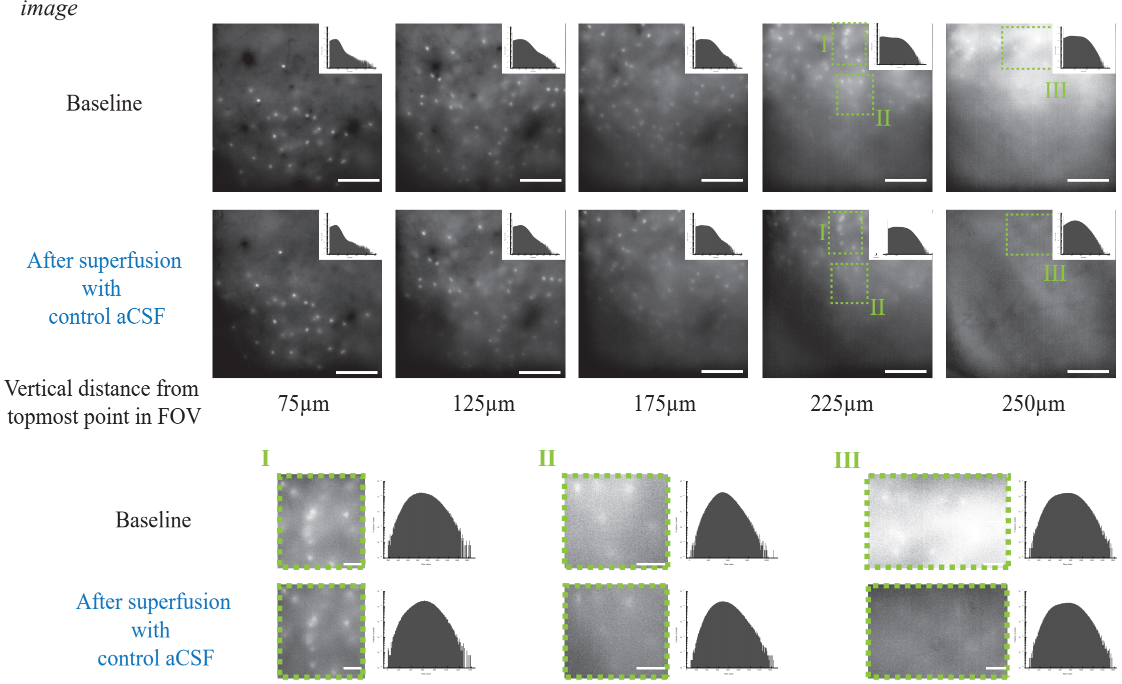


**B**

### DEXTRAN-ACSF

*Display settings autoscaled to the post-superfusion image of each pair and applied to the paired baseline and post-superfusion images.*
*Note that each ROI I-III was autoscaled separately based on the post-superfusion ROI image, with the same settinsg then applied to the baseline image*

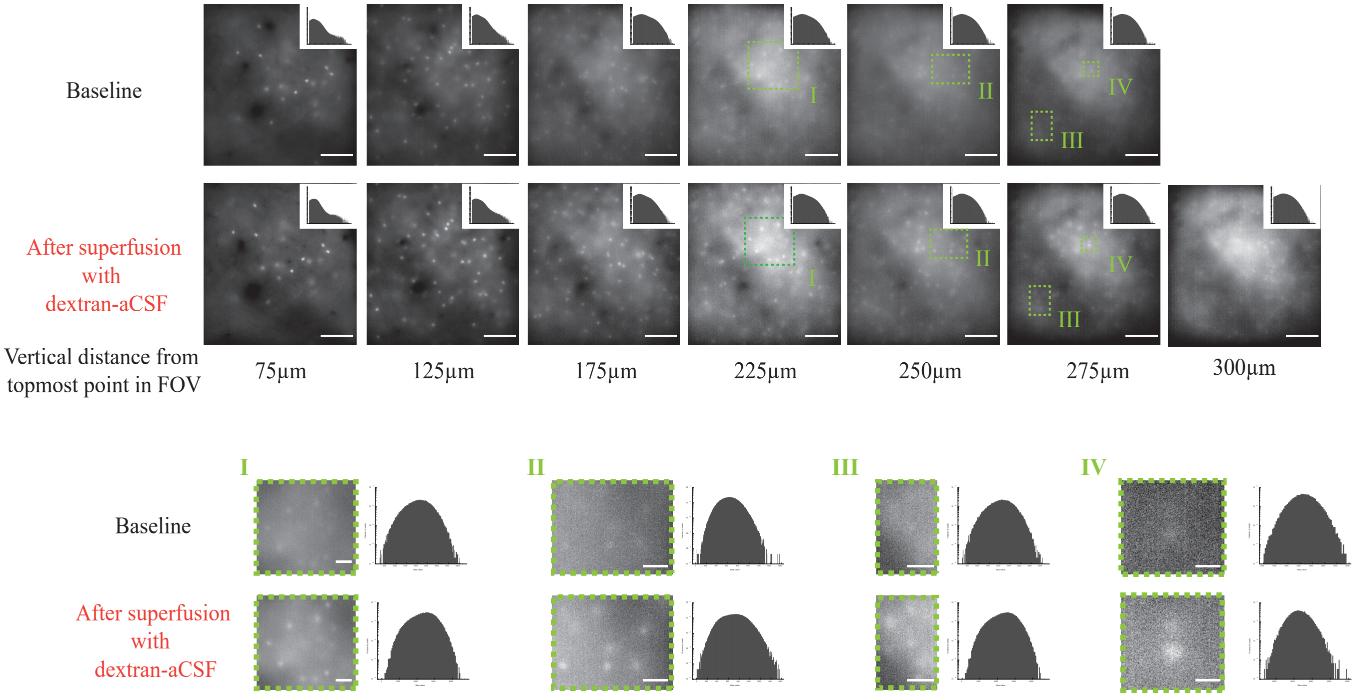

**Fig. S5.**

**Superfusion of the cortical surface for one hour with the modified aCSF enhances imaging at depth under one-photon microscopy.** Same as main text **Figure 2**, with the images displayed using the brightness, contrast, minimum and maximum values determined by autoscaling in Fiji for the post-superfusion image of each pair. See **Methods** for details. **A.** Representative maximum intensity projections, after background subtraction (see **Results** and **Methods** for details), for different depths, of tdTomato-labeled PV+ neurons in mouse somatosensory cortex, before (**top**) and after (**bottom**) 1hr cortical superfusion with control aCSF. Each maximum intensity projection is taken over 7 slices, acquired at 1.5 µm intervals, to compensate for slight misalignments in depth between the two conditions. For both before and after superfusion conditions, the image stacks were acquired under plain aCSF, using identical parameters. The settings for image acquisition and display are the same for the two conditions. Scale bar = 100µm. The highlighted regions of interest (ROI's) **I, II,** and **III** are shown at higher magnification at the bottom of the panel. Scale bar = 25µm. Histograms for the raw pixel values are shown, plotted in semilogarithmic form, in the top right corner of the overall field of view, and to the right of the selected ROIs. The inset in each panel shows the histogram for the raw image, plotted on a semilogarithmic scale. **B.** As in A., for an animal superfused for 1hr with dextran-aCSF instead of control aCSF. For the full-frame images, scale bar = 100µm. For the highlighted regions of interest (ROI's), shown at a larger scale on the right: in **I, II**, and **VI** scale bar = 25µm; in **III** scale bar = 10µm.

**A**

### SUPERFUSION WITH CONTROL ACSF

*Subpanels from Figure 4A, autoscaled in Fiji to maximize the visibility of dim cells, the same display settinsg being then used for both images of a given matching pair.*

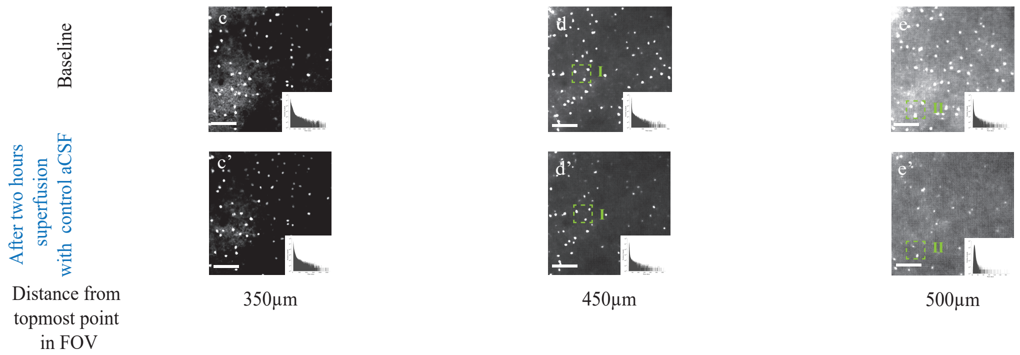


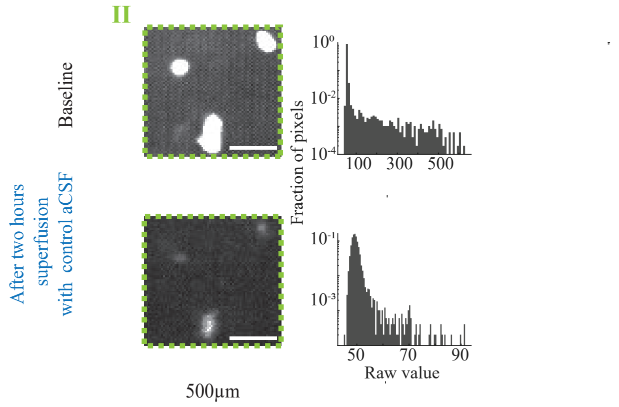


**B**

### SUPERFUSION WITH DEXTRAN ACSF

*Subpanels from Figure 4B, autoscaled in Fiji to maximize the visibility of dim cells, the same display settinsg being then used for both images of a given matching pair.*

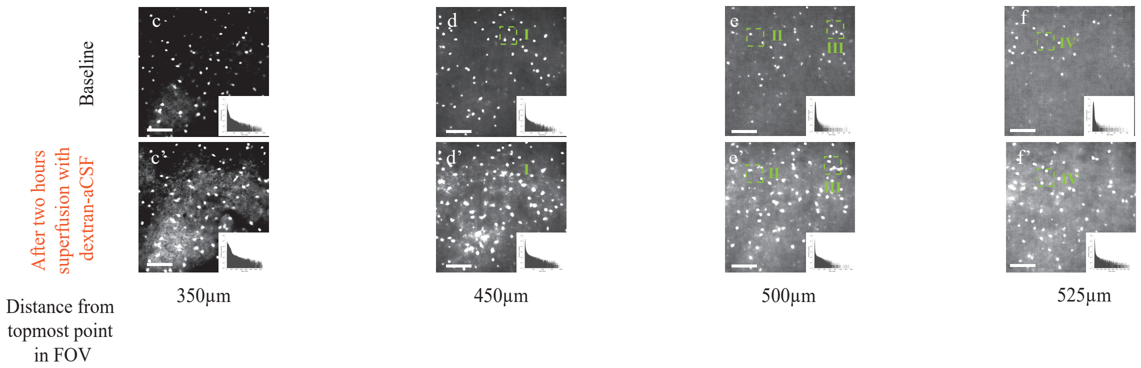


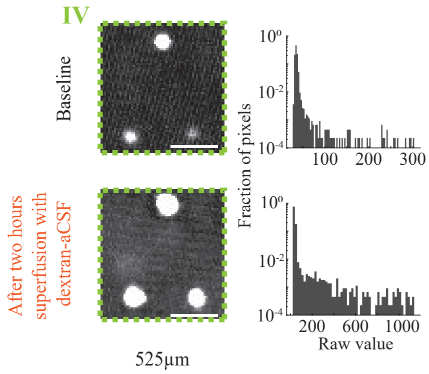

**Fig. S6.**

**Optical clearing at depth observed with two photon imaging in vivo of fluorescently labeled primary visual cortex neurons.** Same as main text **Figure 4**, with the images displayed using the brightness, contrast, minimum and maximum values determined by autoscaling in Fiji for the post-superfusion image of each pair. See **Methods** for details. **A.** Representative average intensity projections, each from 2 imaging slices taken at 2.5μm intervals, from different depths of tdTomato labeled PV+ neurons in mouse primary visual cortex before (top) and after (bottom) 2hrs cortical superfusion with control aCSF. Scale bar = 200μm Top: before superfusion with control aCSF. Bottom: after 2hrs superfusion. Acquisition and display settings are the same for the two conditions. In the top set of images the full field of view is shown, while below them the enlargements of two highlighted ROI's (**I** and **II**) are shown, together with the corresponding histograms of raw pixel values, plotted on a semilogarithmic scale. Scale bar = 25μm. **B.** As in A., for a representative animal superfused with dextran-aCSF. For the image set showing the full field of view, scale bar = 200μm. For ROIs I-IV scale bar = 25μm.

**A**

### SUPERFUSION WITH CONTROL ACSF

*Note that the corresponding baseline and post-superfusion images for each pair are shown with the same display settings, while different pairs of pre- and post-superfusion images use different display settings to balance visibility of dim cells with minimizing oversaturation.*

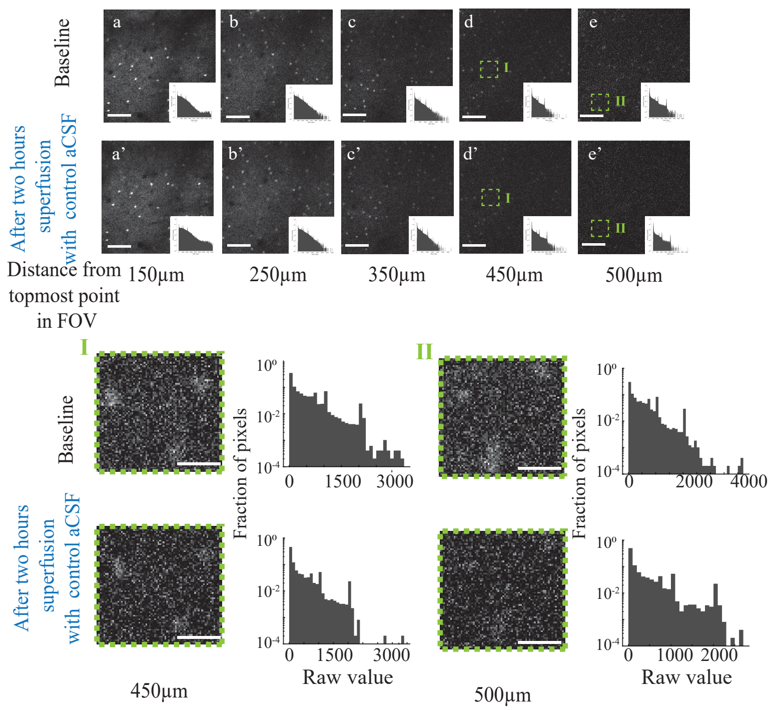


**B**

### SUPERFUSION WITH DEXTRAN ACSF

*Note that the corresponding baseline and post-superfusion images for each pair are shown with the same display settings, while different pairs of pre- and post-superfusion images use different display settings to balance visibility of dim cells with minimizing oversaturation.*

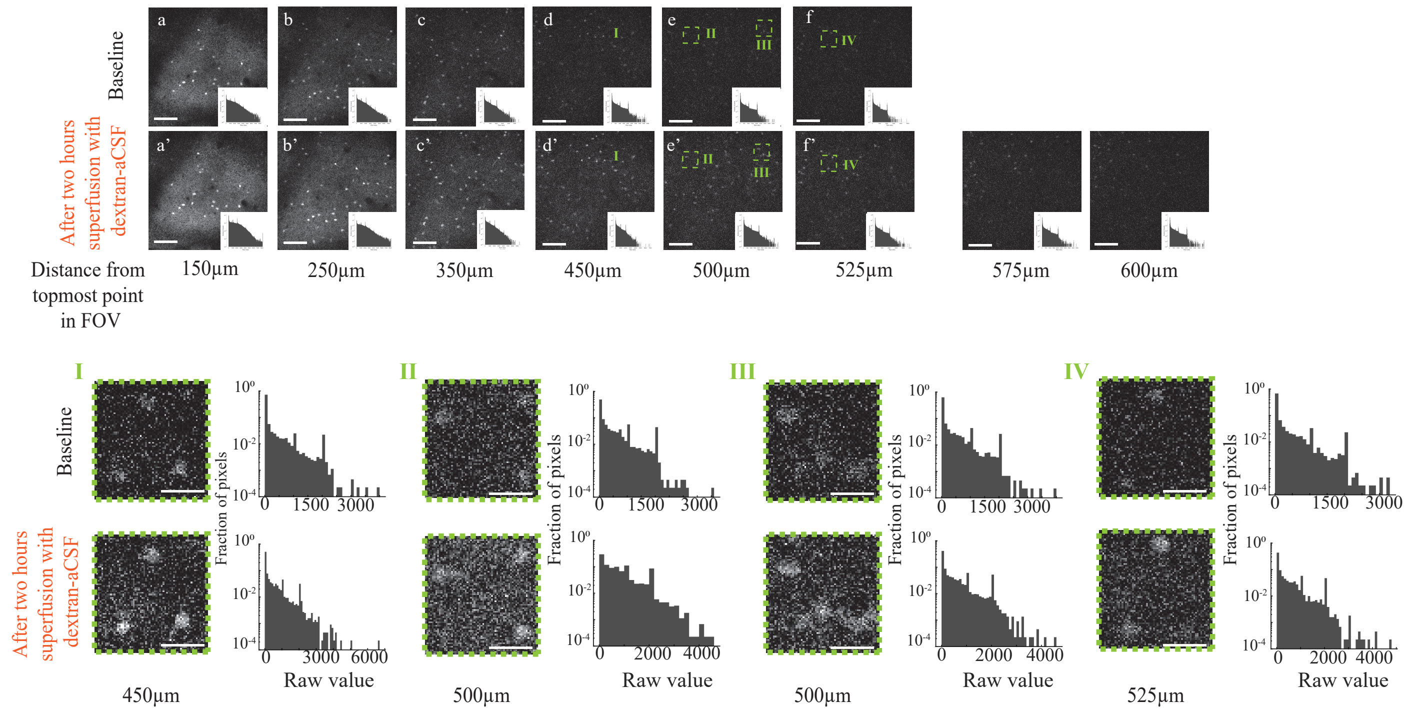

**Fig. S7.**

**As in Figure 4, with unprocessed (i.e. not denoised) images.** For both A and B, the scale bar in the full field of view panels is 150µm, those in the ROIs shown at higher magnification 37.5 µm.

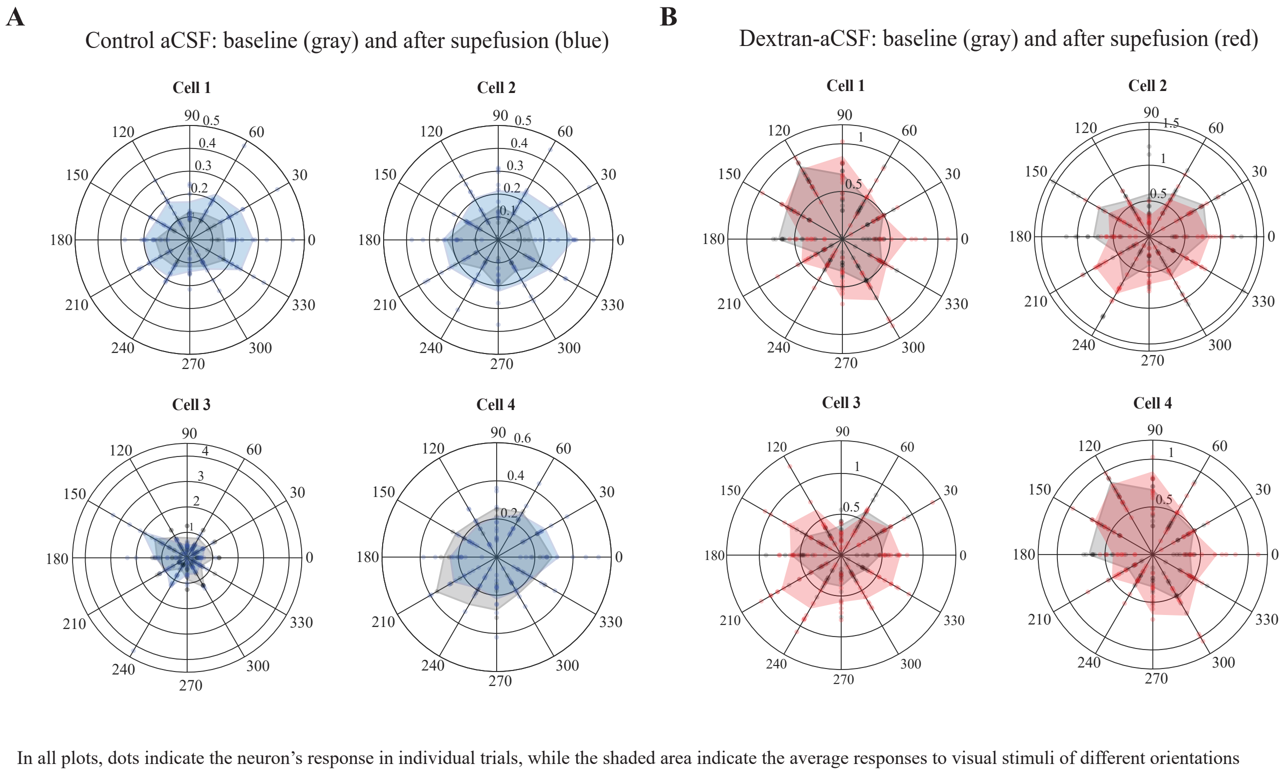


**Fig. S8.**

**Polar plots of individual and trial-averaged responses to visual stimuli of representative cells with baseline OSI or DSI values between 0.1 and 0.2, showing preservation of visual tuning after cortical superfusion for two hours with either control aCSF or dextran-aCSF.** Colors and symbols as in **Fig. 6.**

**A**

Control aCSF: OSI and DSI values at baseline (gray) and after superfusion (blue)

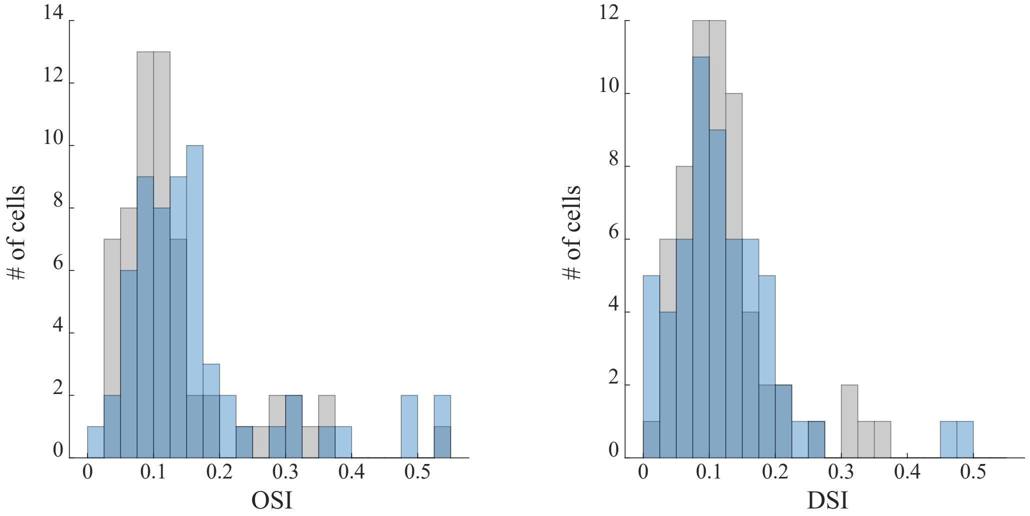


**B**

Dextran aCSF: OSI and DSI values at baseline (gray) and after superfusion (red)

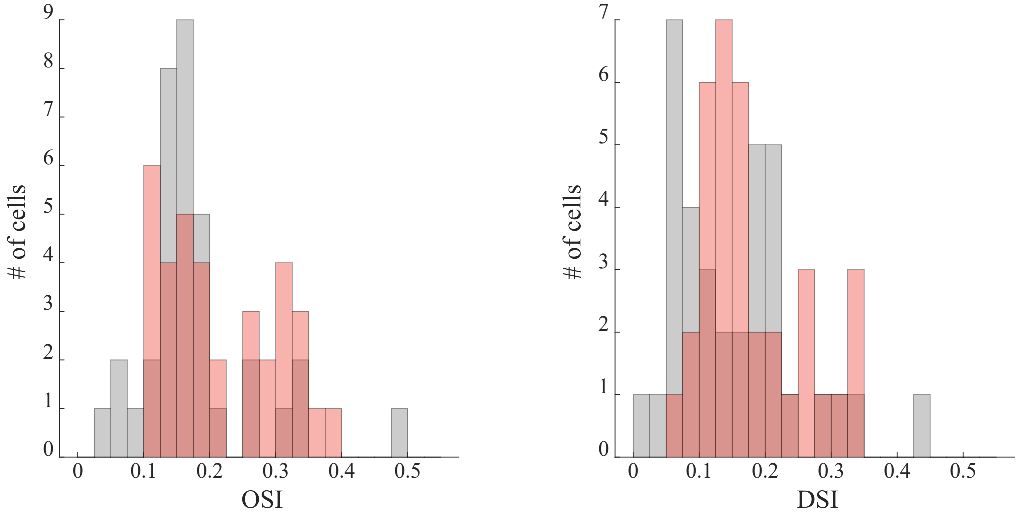


**Fig. S9.**

**Orientation Selectivity Index (OSI) and Direction Selectivity Index (DSI) values for all matched (before and after superfusion) cells.**

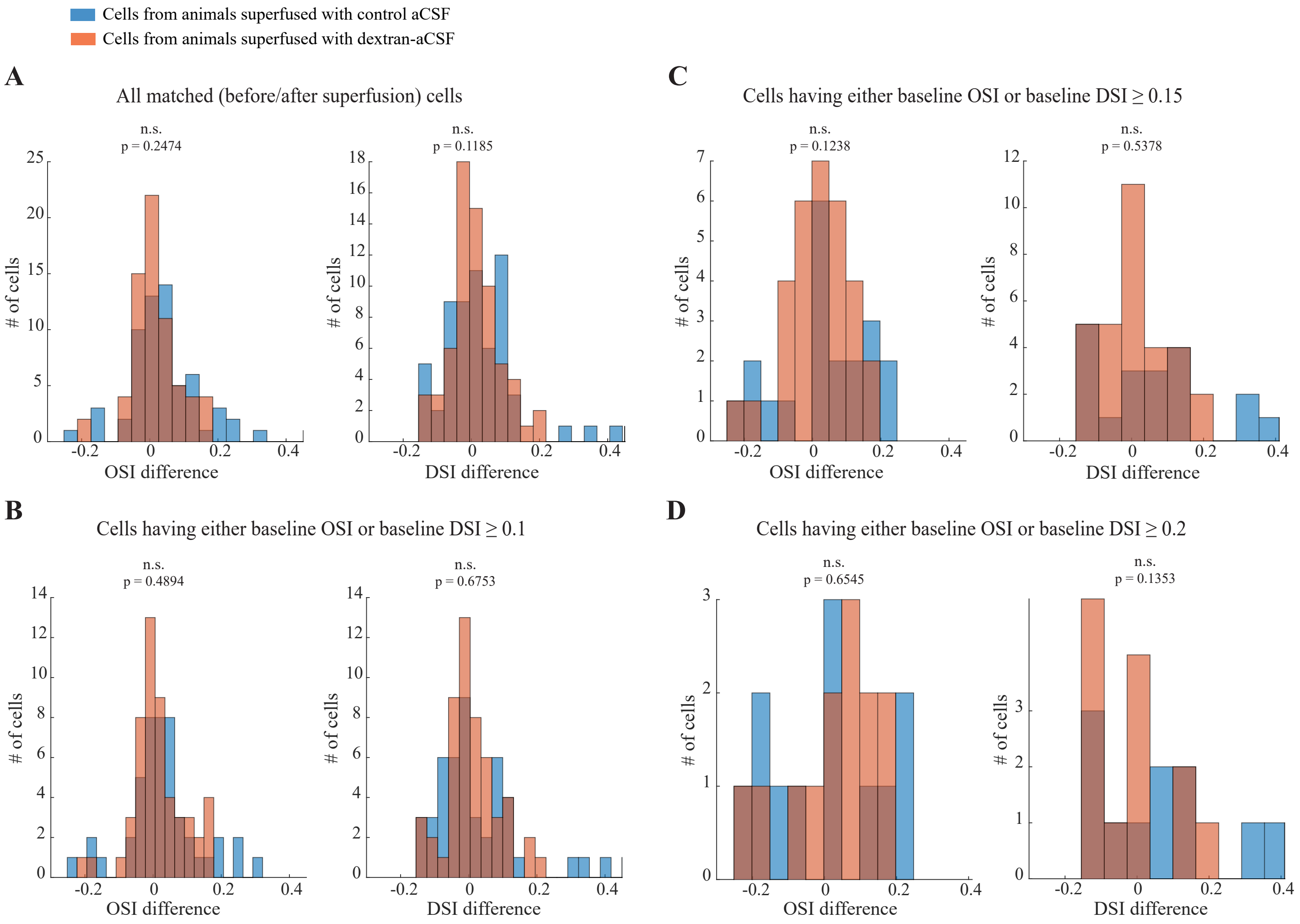


**Fig. S10.**

**Comparison of changes in Orientation Selectivity Index (OSI) and Direction Selectivity Index (DSI) values following two hours superfusion with control aCSF vs. with dextran aCSF.** Reported p-values for two-sample Kolmogorov-Smirnov test. See Results for full statistics.

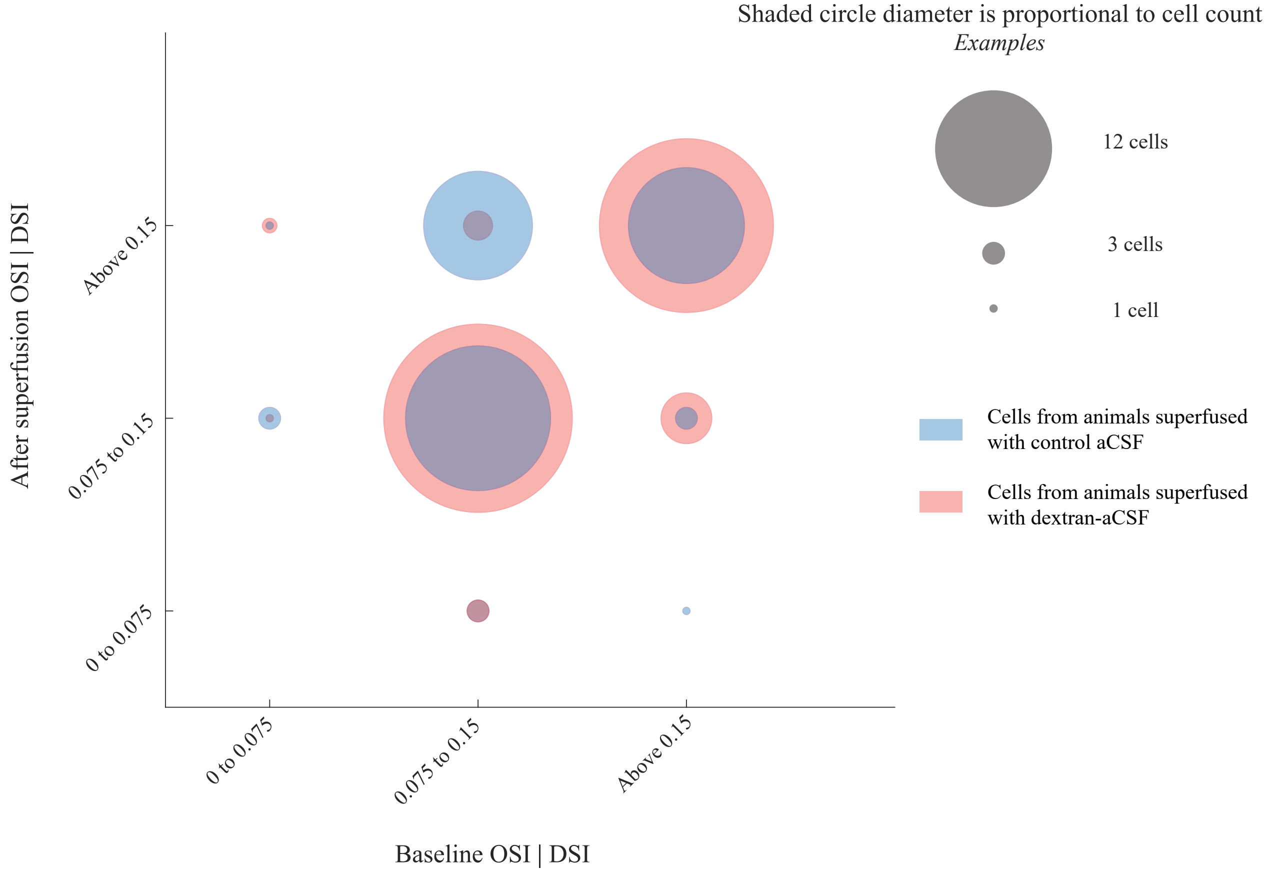


**Fig. S11. OSI and DSI joint changes following superfusion with either control aCSF or dextran-aCSF.** Cells that show visual selectivity at baseline almost always retain it after superfusion and, conversely, cells that don't show visual tuning at baseline rarely show strong visual tuning after superfusion, in both control aCSF and dextran-aCSF superfused animals.

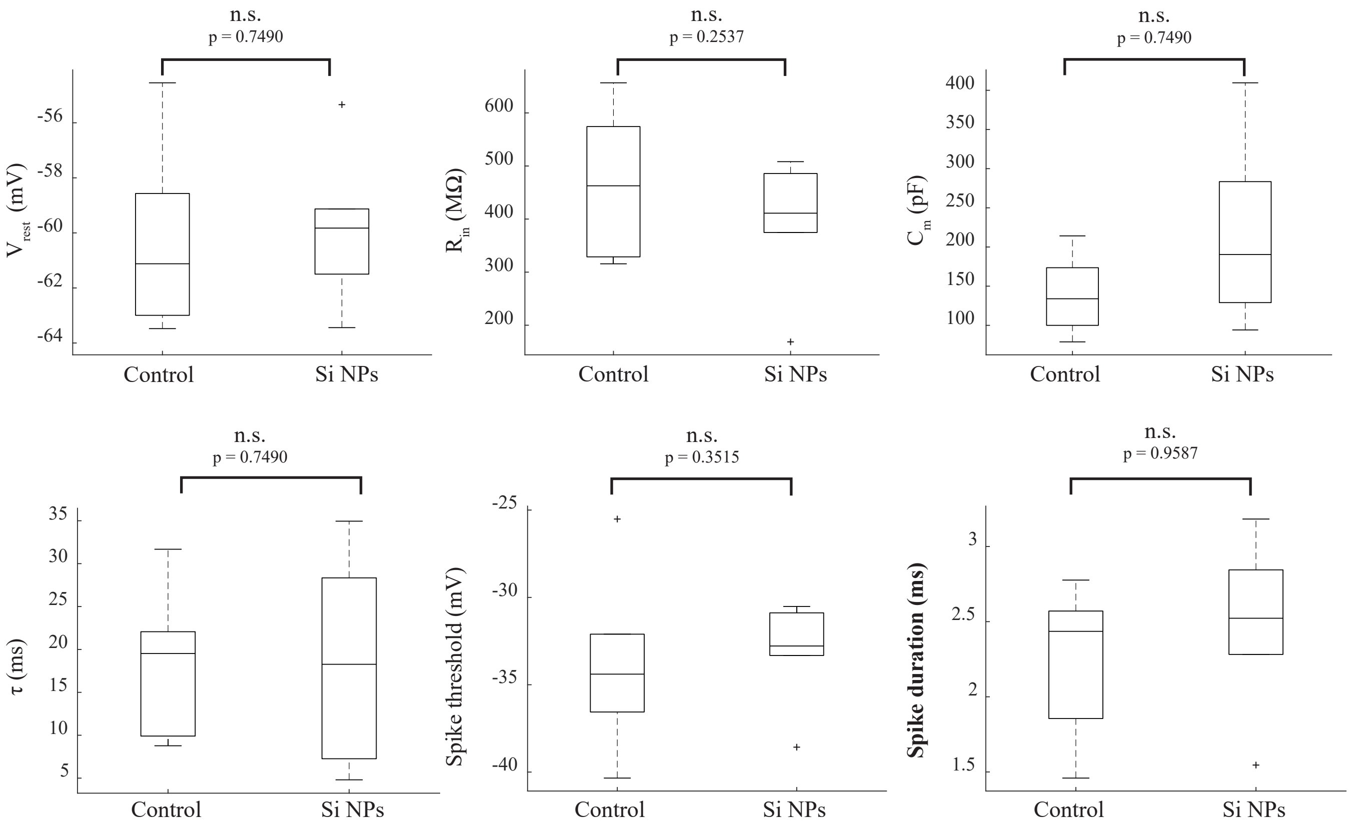


**Fig. S12.**

**Electrophysiological properties of primary neuronal cultures assessed by whole-cell patch clamping after one hour incubation with Tyrode containing a concentration of Si NCs sufficient to raise the refractive index by 0.01.** Electrophysiological properties of primary neuronal cultures assessed by whole-cell patch clamping, comparing control conditions and one hour incubation with Tyrode solution containing the same concentration of Si NCs as the clearing solution used in A. and B. From top left: resting membrane potential, input resistance, membrane capacitance, membrane time constant, spike threshold, and spike duration. Please see Results for full statistics. Throughout the figure, an asterisk denotes statistical significance at the 0.05 level, n.s. = not significant.

| Change in bead brightness above background (%) | | Control aCSF | Clearing aCSF, with R.I. 0.01 greater than control aCSF | | |
|---|---|---|---|---|---|
| | | | Dextran-aCSF | PEG-aCSF | Iodixanol-aCSF |
| | 20th percentile | -64.18 | +191.16 | +297.94 | +71.36% |
| | Mean | -55.85 | +249.14 | +521.42 | +124.39 |
| | Median | -60.18 | +217.67 | +429.21 | +127.63 |
| | 80th percentile | -45.20 | +271.67 | +818.67 | +200.36 |
| Number of beads at baseline | | 51 | 67 | 82 | 90 |
| Number of beads after incubation | | 46 | 69 | 124 | 244 |
| Number of slices | | 5 | 5 | 4 | 7 |
| Number of mice | | 2 | 3 | 3 | 3 |
| p (post-incubation vs. baseline) | | 0.0201 | $6.4582*10^{-29}$ | $4.4210*10^{-41}$ | $4.3646*10^{-6}$ |
| p (clearing aCSF vs. control aCSF) | | N/A | $2.9791*10^{-26}$ | 0.0069 | 0.0016 |

**Table S1. Change in signal intensity above background from beads imaged through acute brain slices after incubation for one hour with control aCSF or aCSF with refractive index increased by 0.01 by the addition of either 1.5mM Dextran 40kDa, 6mM PEG 10kDa, or 40mM iodixanol.** Reported p-values for both post-incubation vs. baseline and clearing-aCSF vs. control aCSF comparisons were calculated using the default MATLAB (Mathworks, NJ) implementation of the two-sample Kolmogorov-Smirnov test.

| | | Vertical distance from topmost point of the brain surface within the field of view | | | | | |
|---|---|---|---|---|---|---|---|
| | | ≤ 125µm | | >125µm | | Aggregate across all distances | |
| Change in soma brightness above background (%) | | Control-aCSF | Dextran-aCSF | Control-aCSF | Dextran-aCSF | Control-aCSF | Dextran-aCSF |
| | $20^{th}$ percentile | -40.68 | +43.23 | -44.84 | +66.79 | -44.50 | +56.56 |
| | Mean | -12.82 | +146.33 | -13.51 | +156.25 | -13.51 | +153.08 |
| | Median | -8.19 | +74.48 | -18.48 | +110.59 | -14.82 | +99.22 |
| | $80^{th}$ percentile | +12.35 | +133.40 | +15.93 | +185.96 | +15.20 | +169.24 |
| Number of cells | | 198 | 411 | 605 | 874 | 803 | 1285 |
| Number of animals | | 3 | 4 | 3 | 4 | 3 | 4 |
| p (post-superfusion vs. baseline) | | $2.0276*10^{-7}$ | $3.6842*10^{-68}$ | $1.4976*10^{-27}$ | $3.22793*10^{-144}$ | $1.359810^{-31}$ | $2.5189*10^{-208}$ |
| p (≤ 125µm vs. >125µm) | | N/A | N/A | $4.0652*10^{-4}$ | $1.9289*10^{-17}$ | N/A | N/A |
| p (dextran-aCSF vs. control aCSF) | | N/A | $2.2619*10^{-83}$ | N/A | $6.7683*10^{-245}$ | N/A | $1.3419*10^{-319}$ |

**Table S2. Effect of one hour superfusion with dextran-aCSF or control solution on the signal above background from fluorescently labeled neurons in mouse primary sensory cortex in vivo, imaged under one-photon microscopy.** Reported p-values for both ≤ 125µm vs. > 125µm distances and dextran-aCSF vs. control aCSF comparisons were calculated using the default MATLAB (Mathworks, NJ) implementation of the two-sample Kolmogorov-Smirnov test. For comparing the brightness above background of the same cells before and after incubation, using the default MATLAB (Mathworks, NJ) implementation of the Wilcoxon signed rank test was used instead.

| | Vertical distance from topmost point of the brain surface within the field of view | | | | | | | | |
|---|---|---|---|---|---|---|---|---|---|
| | ≤ 125µm | | | >125 µm | | | Aggregate across all distances | | |
| **Change in soma brightness above background (%)** | Control-aCSF | PEG-aCSF | Iodixanol-aCSF | Control-aCSF | PEG-aCSF | Iodixanol-aCSF | Control-aCSF | PEG-aCSF | Iodixanol-aCSF |
| **20$^{th}$ percentile** | -28.76 | +15.57 | +13.29 | -46.27 | +35.17 | +16.14 | -44.56 | +32.94 | +15.53 |
| **Mean** | -10.89 | +78.80 | +85.90 | -13.45 | +103.73 | +168.23 | -13.17 | +100.5 | +119.54 |
| **Median** | -5.09 | +71.73 | +51.40 | -16.75 | +68.95 | +123.07 | -14.82 | +69.45 | +62.46 |
| **80$^{th}$ percentile** | +10.46 | 123.44 | +106.56 | +16.11 | +154.99 | +227.76 | +15.31 | +147.24 | +181.92 |
| **Number of cells** | 87 | 49 | 139 | 718 | 283 | 96 | 805 | 332 | 235 |
| **Number of animals** | 3 | 6 | 9 | 3 | 6 | 9 | 3 | 6 | 9 |
| **p (post-superfusion vs. baseline)** | 0.0014 | $5.0951 *10^{-9}$ | $1.6353 *10^{-17}$ | $1.1505 *10^{-29}$ | $1.3536 *10^{-47}$ | $3.8391 *10^{-13}$ | $2.1980 *10^{-31}$ | $2.1086 *10^{-65}$ | $4.0585 *10^{-34}$ |
| **p (≤ 125µm vs. >125µm)** | N/A | N/A | N/A | $7.5688*10^{-5}$ | 0.123 | $1.2022 *10^{-6}$ | N/A | N/A | N/A |
| **p (clearing-aCSF vs. control aCSF)** | N/A | 0.0140 | $1.4943 *10^{-23}$ | N/A | $3.8100 *10^{-99}$ | $1.5120 *10^{-34}$ | N/A | $1.5690 *10^{-116}$ | $3.5336 *10^{-66}$ |

**Table S3: Effect of one hour superfusion with control aCSF, PEG-aCSF, or iodixanol-aCSF solution on the signal above background from fluorescently labeled neurons in mouse primary sensory cortex in vivo, imaged under one-photon microscopy.** Reported p-values for both ≤ 125µm vs. > 125µm distances and clearing-aCSF vs. control aCSF comparisons were calculated using the default MATLAB (Mathworks, NJ) implementation of the two-sample Kolmogorov-Smirnov test. For comparing the brightness above background of the same cells before and after incubation, using the default MATLAB (Mathworks, NJ) implementation of the Wilcoxon signed rank test was used instead.

| | Vertical distance from topmost point of the brain surface within the field of view | | | | | | | | | |
|---|---|---|---|---|---|---|---|---|---|---|
| | ≤ 250µm | | 250-350 µm | | 350-450 µm | | 450-550 µm | | Aggregate across all distances | |
| **Change in soma brightness above background (%)** | Control aCSF | Dextran-aCSF | Control aCSF | Dextran-aCSF | Control aCSF | Dextran-aCSF | Control aCSF | Dextran-aCSF | Control aCSF | Dextran-aCSF |
| **20th percentile** | -25.60 | 28.18 | -43.24 | 43.11945 | -77.54 | 77.18 | -88.24 | 50.16 | -58.40 | 43.21 |
| **Median** | -3.80 | 60.09 | -17.60 | 90.86 | -57.19 | 204.31 | -79.03 | 338.42 | -23.04296 | 97.09 |
| **80th percentile** | 32.17 | 93.01 | 54.97 | 152.10 | -20.22 | 376.91 | -52.49 | 1042.62 | 33.48 | 254.34 |
| **Number of cells** | 322 | 648 | 343 | 697 | 285 | 728 | 42 | 243 | 992 | 2316 |
| **Number of animals** | 2 | 3 | 2 | 3 | 2 | 3 | 2 | 3 | 2 | 3 |
| **p (post-superfusion vs. baseline)** | $1.42*10^{-6}$ | 0* | $1.91*10^{-14}$ | 0* | $3.26*10^{-40}$ | 0* | $1.10953*10^{-7}$ | $1.49*10^{-38}$ | 0* | 0* |
| **p (dextran-aCSF vs. control aCSF)** | N/A | $2.59*10^{-66}$ | N/A | $5.65*10^{-67}$ | N/A | $7.13*10^{-118}$ | N/A | $5.14*10^{-26}$ | N/A | $4.86*10^{-257}$ |

**Table S4: Effect of two hours superfusion with dextran-aCSF or control solution on the signal above background from fluorescently labeled neurons in mouse primary sensory cortex in vivo, imaged under two-photon microscopy.** Reported p-values for dextran-aCSF vs. control aCSF comparisons were calculated using the default MATLAB (Mathworks, NJ) implementation of the two-sample Kolmogorov-Smirnov test. For comparing the brightness above background of the same cells before and after incubation the default MATLAB (Mathworks, NJ) implementation of the Wilcoxon signed rank test was used instead.

p = 0* indicates that the number was small enough to be rounded off to 0 by the default MATLAB implementation of the Wilcoxon signed-rank test.

| Comparison of changes in soma brightness above background following superfusion with control aCSF | | | | | | |
|---|---|---|---|---|---|---|
| p-value (two-sample Kolmogorov-Smirnov test) | | Vertical distance from topmost point of the brain surface within the field of view | | | | |
| | | **≤ 250µm** | **250-350 µm** | **350-450 µm** | **450-550 µm** | **Aggregate across all distances** |
| **Vertical distance from topmost point of the brain surface within the field of view** | **≤ 250µm** | N/A | 3.73E-09 | 5.32E-56 | 1.53E-22 | 4.12E-19 |
| | **250-350 µm** | 3.73E-09 | N/A | 1.36E-32 | 4.27E-18 | 5.67E-05 |
| | **350-450 µm** | 5.32E-56 | 1.36E-32 | N/A | 8.83E-07 | 1.89E-25 |
| | **450-550 µm** | 1.53E-22 | 4.27E-18 | 8.83E-07 | N/A | 1.63E-13 |
| **N (cells)** | | 322 | 343 | 285 | 42 | 992 |
| **N (mice)** | | 2 | 2 | 2 | 2 | 2 |

| Comparison of changes in soma brightness above background following superfusion with dextran-aCSF | | | | | | |
|---|---|---|---|---|---|---|
| p-value (two-sample Kolmogorov-Smirnov test) | | Vertical distance from topmost point of the brain surface within the field of view | | | | |
| | | **≤ 250µm** | **250-350 µm** | **350-450 µm** | **450-550 µm** | **Aggregate across all distances** |
| **Vertical distance from topmost point of the brain surface within the field of view** | **≤ 250µm** | N/A | 9.55E-32 | 1.46E-123 | 5.81E-59 | 1.33E-50 |
| | **250-350 µm** | 9.55E-32 | N/A | 7.80E-62 | 5.17E-45 | 9.81E-12 |
| | **350-450 µm** | 1.46E-123 | 7.80E-62 | N/A | 1.57E-13 | 2.48E-46 |
| | **450-550 µm** | 5.81E-59 | 5.17E-45 | 1.57E-13 | N/A | 1.32E-28 |
| **N (cells)** | | 648 | 697 | 728 | 243 | 2316 |
| **N (mice)** | | 3 | 3 | 3 | 3 | 3 |

**Table S5: Statistical comparison of the effect for different depth ranges of two hours superfusion with dextran-aCSF or control solution on the signal above background from fluorescently labeled neurons in mouse primary sensory cortex in vivo, imaged under two-photon microscopy.**

| | | Number of cells in a given range of DSI or OSI values (considering the higher of the two indices) after superfusion with control aCSF | | | | | |
|---|---|---|---|---|---|---|---|
| Number of cells in a given range of DSI or OSI values (considering the higher of the two indices) before superfusion with control aCSF | | **x ≤ 0.05** | **0.05 < x ≤ 0.1** | **0.1 < x ≤ 0.15** | **0.15 < x ≤ 0.2** | **x > 0.2** | **Total** |
| | **x ≤ 0.05** | 0 | 0 | 0 | 0 | 0 | 0 |
| | **0.05 < x ≤ 0.1** | 1 | 2 | 5 | 4 | 3 | 15 |
| | **0.1 < x ≤ 0.15** | 0 | 7 | 11 | 7 | 2 | 27 |
| | **0.15 < x ≤ 0.2** | 0 | 1 | 0 | 3 | 4 | 8 |
| | **x > 0.2** | 0 | 2 | 1 | 1 | 8 | 12 |
| | **Total** | 1 | 12 | 17 | 15 | 17 | 62 |

| | | Number of cells in a given range of DSI or OSI values (considering the higher of the two indices) after superfusion with dextran-aCSF | | | | | |
|---|---|---|---|---|---|---|---|
| Number of cells in a given range of DSI or OSI values (considering the higher of the two indices) superfusion with dextran-aCSF | | **x ≤ 0.05** | **0.05 < x ≤ 0.1** | **0.1 < x ≤ 0.15** | **0.15 < x ≤ 0.2** | **x > 0.2** | **Total** |
| | **x ≤ 0.05** | 0 | 0 | 0 | 0 | 0 | 1 |
| | **0.05 < x ≤ 0.1** | 1 | 2 | 5 | 4 | 3 | 14 |
| | **0.1 < x ≤ 0.15** | 0 | 7 | 11 | 7 | 2 | 21 |
| | **0.15 < x ≤ 0.2** | 0 | 1 | 0 | 3 | 4 | 18 |
| | **x > 0.2** | 0 | 2 | 1 | 1 | 8 | 13 |
| | **Total** | 0 | 20 | 17 | 10 | 20 | 67 |

**Table S6. Number of cells with either DSI or OSI values (considering the higher of the two indices) in a given range before and after superfusion with either control aCSF or dextran-aCSF – Selectivity Index ranges: below 0.05, between 0.05 and 0.1, between 0.1 and 0.15, between 0.15 and 0.2, and above 0.2.**

| | Number of cells in a given range of DSI or OSI values (considering the higher of the two indices) before superfusion with control aCSF | | | | |
|---|---|---|---|---|---|
| Number of cells in a given range of DSI or OSI values (considering the higher of the two indices) after superfusion with control aCSF | | x ≤ 0.075 | 0.075 < x ≤ 0.15 | x > 0.15 | Total |
| | x ≤ 0.075 | 0 | 3 | 1 | 4 |
| | 0.075 < x ≤ 0.15 | 3 | 20 | 15 | 38 |
| | x > 0.15 | 1 | 3 | 16 | 20 |
| | Total | 4 | 26 | 32 | 62 |

| | Number of cells in a given range of DSI or OSI values (considering the higher of the two indices) before superfusion with dextran-aCSF | | | | |
|---|---|---|---|---|---|
| Number of cells in a given range of DSI or OSI values (considering the higher of the two indices) after superfusion with dextran-aCSF | | x ≤ 0.075 | 0.075 < x ≤ 0.15 | x > 0.15 | Total |
| | x ≤ 0.075 | 0 | 1 | 2 | 3 |
| | 0.075 < x ≤ 0.15 | 3 | 26 | 4 | 33 |
| | x > 0.15 | 0 | 7 | 24 | 31 |
| | Total | 3 | 34 | 30 | 67 |

**Table S7. Number of cells with either DSI or OSI values (considering the higher of the two indices) in a given range before and after superfusion with either control aCSF or dextran-aCSF – Selectivity Index ranges: below 0.075, between 0.075 and 0.15, and above 0.15**

| Change in soma brightness above background (%) | | Control-aCSF | Dextran-aCSF |
|---|---|---|---|
| | 20th percentile | +15.3 | +48.15 |
| | Mean | +29.9 | +190.60 |
| | Median | +30.4 | +183.23 |
| | 80th percentile | +40.9 | +23.20 |
| **Number of cells** | | 81 | 73 |
| **Number of animals** | | 2 | 3 |
| **p (post-superfusion vs. baseline)** | | $2.3231*10^{-10}$ | $4.5315*10^{-20}$ |
| **p (dextran-aCSF vs. control aCSF)** | | N/A | $1.8553*10^{-45}$ |

**Table S8. Effect of two hours superfusion with control aCSF or dextran-aCSF on the signal above background recorded in vivo under two-photon microscopy from primary visual cortex neurons that expressed a genetically encoded calcium indicator.** Reported p-values for dextran-aCSF vs. control aCSF comparisons were calculated using the default MATLAB (Mathworks, NJ) implementation of the two-sample Kolmogorov-Smirnov test. For comparing the brightness above background of the same cells before and after incubation, using the default MATLAB (Mathworks, NJ) implementation of the Wilcoxon signed rank test was used instead.

| | | Iodixanol-aCSF | |
|---|---|---|---|
| Change in soma brightness above background (%) | | **Green channel** (475/28 nm excitation, emission 535/22 band pass filter) | **Red channel** (excitation 637nm, emission 664nm long pass filter) |
| | 20th percentile | 14.38 | +37.35 |
| | Mean | 83.82 | +136.57 |
| | Median | 60.26 | +110.18 |
| | 80th percentile | 131.18 | +221.67 |
| Number of cells | | 37 | 37 |
| Number of areas within the slices | | 11 | 11 |
| Number of slices | | 9 | 9 |
| Number of animals | | 4 | 4 |
| p (post-superfusion vs. baseline) | | $1.2569*10^{-5}$ | $1.8650*10^{-7}$ |

**Table S9. Effect of one hour superfusion with carbogenated iodixanol-aCSF on the signal above background recorded in acute brain slices from neurons expressing the genetically encoded voltage indicator Archon-GFP.** Reported p-values for comparing the brightness above background of the same cells before and after incubation were calculated using the default MATLAB (Mathworks, NJ) implementation of the Wilcoxon signed rank test.

| Change in bead brightness above background (%) | | Control aCSF | Si NPs-aCSF |
|---|---|---|---|
| | **20th percentile** | -64.18 | +110.96 |
| | **Mean** | -55.85 | +300.74 |
| | **Median** | -60.18 | +171.94 |
| | **80th percentile** | -45.20 | +580.68 |
| **Number of beads at baseline** | | 51 | 40 |
| **Number of beads after incubation** | | 46 | 69 |
| **Number of slices** | | 5 | 5 |
| **Number of mice** | | 2 | 2 |
| **p (post-incubation vs. baseline)** | | 0.0201 | $5.2458*10^{-8}$ |
| **p (clearing aCSF vs. control aCSF)** | | N/A | 0.0069 |

**Table S10. Change in signal intensity above background from beads imaged through acute brain slices after incubation for one hour with control aCSF or aCSF with refractive index increased by 0.01 by the addition of custom PEG-ylated silicon nanoparticles.** Reported p-values for both post-incubation vs. baseline and clearing-aCSF vs. control aCSF comparisons were calculated using the default MATLAB (Mathworks, NJ) implementation of the two-sample Kolmogorov-Smirnov test.
